# StrainCascade: An automated, modular workflow for high-throughput long-read bacterial genome reconstruction and characterization

**DOI:** 10.64898/2026.02.04.698786

**Authors:** Sebastian B.U. Jordi, Isabel Baertschi, Jiaqi Li, Nadia Fasel, Benjamin Misselwitz, Bahtiyar Yilmaz

## Abstract

Long-read sequencing offers unprecedented opportunities for high-resolution bacterial genome reconstruction, yet fragmented bioinformatics workflows hinder biological insights. *StrainCascade* addresses this gap by providing a fully automated, modular pipeline that integrates genome assembly, accurate annotation, and comprehensive functional profiling into a single, reproducible framework. By integrating deterministic execution strategies with a systematic resolution of strain-level structural and functional variability, *StrainCascade* provides high-resolution comparative genomics of strain diversity, host-microbe interactions, antimicrobial resistance, and mobile genetic element dynamics.

## INTRODUCTION

Decoding microbial genomes is foundational to understanding how bacteria adapt, survive, and interact with their environments. From shaping nutrient cycles in ecosystems to modulating inflammation in the human gut^1^, microbial functions are inextricably linked to their genetic makeup. High-resolution genome analysis has become indispensable not only for exploring microbial diversity and evolution but also for pinpointing traits that underlie pathogenicity, antimicrobial resistance, and host-microbe interactions.

Bacterial whole genome sequencing (WGS) has emerged as the key strategy for translating this genomic potential into actionable insights, providing the resolution required for outbreak tracking, pathogen surveillance, and comparative functional genomics. While short-read sequencing platforms, such as Illumina, have long dominated bacterial genome sequencing due to their high accuracy, their inability to resolve repetitive regions, structural variations, and plasmid content remains a major limitation^2^. In contrast, long-read sequencing technologies from Pacific Biosciences (PacBio) and Oxford Nanopore Technologies (ONT) have emerged as powerful alternatives, offering superior contiguity, structural accuracy, and the ability to assemble complete bacterial genomes *de novo*. Although early long-read platforms were hindered by high error rates (∼10-15%), modern PacBio HiFi and ONT R10.4 chemistries now achieve significantly improved accuracy (∼99.9%)^7^. Despite these technical advances, the specialized bioinformatics required to process long-read data continues to limit broad adoption across laboratories.

The transition from short-read to long-read sequencing presents distinct bioinformatics challenges. Short-read WGS benefits from established pipelines such as *TORMES*^3^, *Bactopia*^4^, *ASA*^3^ *P*^5^, and *EnteroBase*^6^ which provide streamlined workflows. In contrast, long-read sequencing requires extra specialized computational tools and many existing bioinformatics pipelines remain optimized for short-read data, lacking the flexibility to efficiently process long-read outputs. These limitations become particularly evident in multi-genome projects, where researchers face three critical challenges: (i) the absence of cohesive end-to-end workflows for read processing, assembly, taxonomic classification and functional profiling; (ii) the inability to incorporate species-specific knowledge into standard pipelines; and (iii) output formats poorly suited for downstream comparative or functional analyses, such as pan-genome reconstruction or phylogenetic visualization.

To address these limitations, we developed *StrainCascade*, a fully automated and modular pipeline that streamlines long-read bacterial genome reconstruction and functional characterization. The pipeline integrates all key analytical steps from read pre-processing and genome assembly to taxonomic classification, genome annotation, and in-depth functional profiling within a unified, reproducible framework using a single-command command-line interface (CLI). Designed to be platform-agnostic, *StrainCascade* automatically optimizes its assembly strategy for both PacBio and ONT reads and supports flexible input formats, including raw reads and pre-assembled contigs. The modular structure allows for customizable execution and re-usability, and optional deterministic processing ensures bit-identical outputs across runs. To demonstrate its performance, we applied *StrainCascade* to a phylogenetically diverse dataset of 152 bacterial genomes (Figure 1), including both public reference genomes and in-house isolates representing diverse human-, animal-, and environmental-associated bacteria. This comprehensive evaluation highlights the capacity of the pipeline to deliver accurate genome reconstruction, high-resolution functional annotation, and lineage-specific insights into genome plasticity and adaptation.

**Figure 1.**
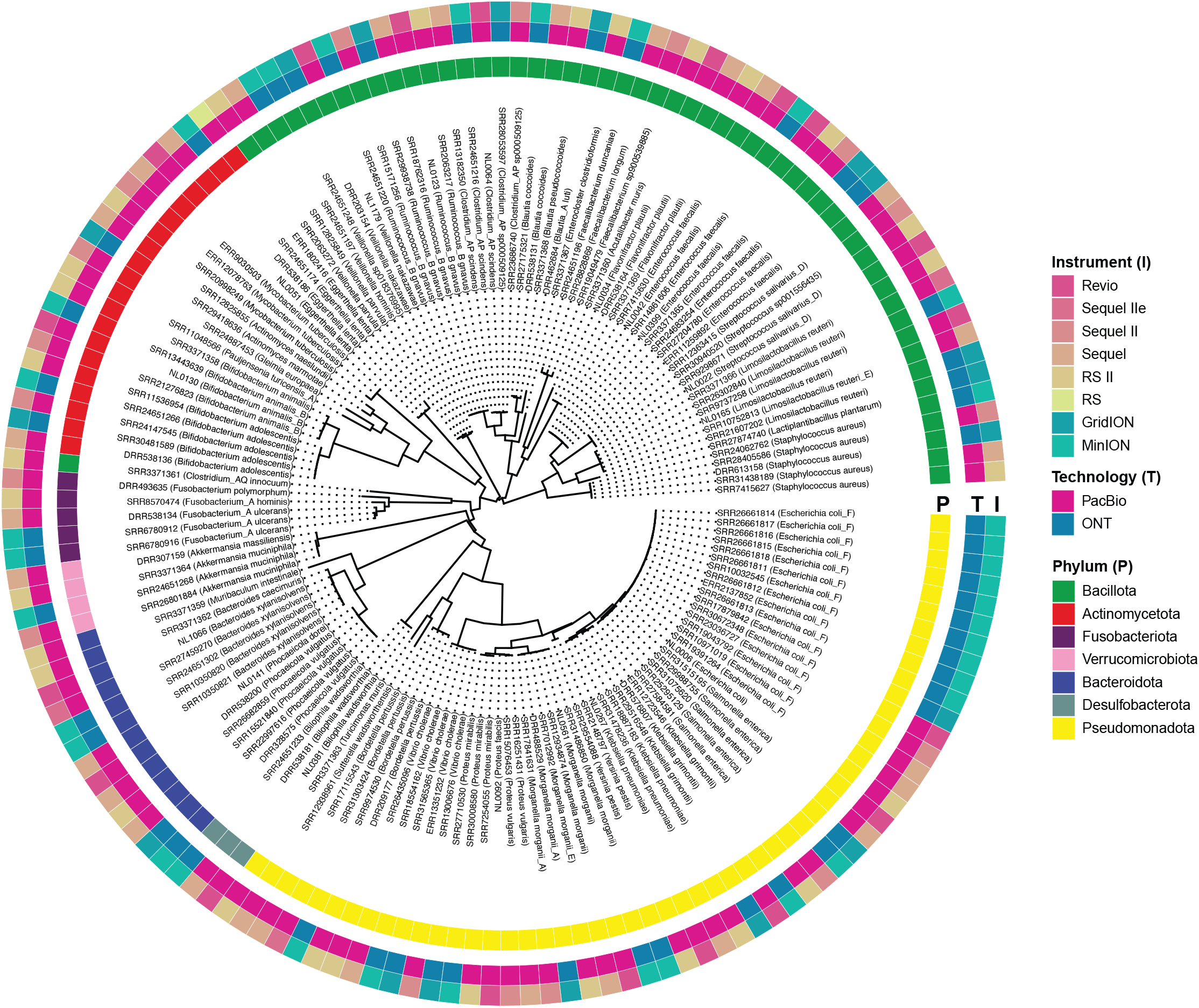
Phylogenetic distribution and sequencing technology metadata of bacterial genomes analyzed with *StrainCascade*. 152 bacterial genomes were used for benchmarking. A phylogenetic tree was constructed using GTDB-Tk *de novo* analysis (de_novo_wf) and implemented in *StrainCascade*. Each branch represents a genome labeled with its closest taxonomic relative, spanning multiple phyla (Bacillota, Actinomycetota, Bacteroidota, Fusobacteriota, Verrucomicrobiota, and Pseudomonadota). Sequencing metadata annotations distinguish PacBio (Revio, Sequel, Sequel II, RS II, RS) and Oxford Nanopore (MinION, GridION) platforms. The dataset integrates genomes from NCBI and in-house patient-derived isolates, ensuring phylogenetic diversity for cross-platform workflow evaluation.

## RESULTS

### Modular architecture and core functionalities of the *StrainCascade* pipeline

*StrainCascade* is organized into a modular, end-to-end framework that automates long-read bacterial genome analysis from raw sequencing data to high-resolution functional insights through a single-command CLI on Unix-based systems. The pipeline is implemented primarily in Shell (57.0%) and R (36.6%), with additional components in Python (6.4%). *StrainCascade* contains 27 sequential modules (SC1–SC27) that together provide a comprehensive, modular framework. The pipeline accepts input in FASTQ, FASTA, or BAM format and supports pre-assembled genomes as input, enabling flexible and customizable workflows with defined logical dependencies (Figure 2, See STAR Methods).

**Figure 2.**
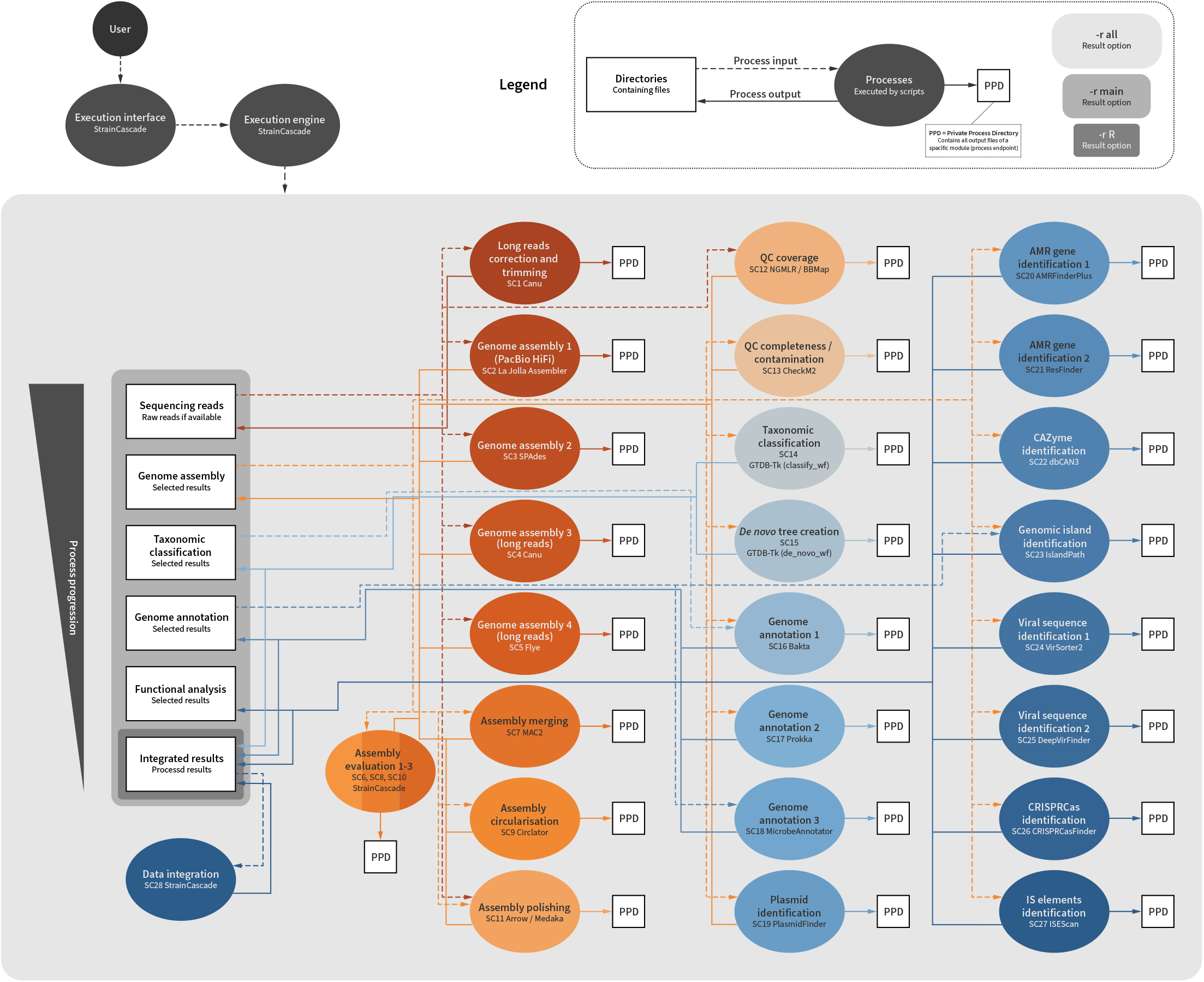
Modular workflow of *StrainCascade* for high-throughput bacterial genome analysis. The pipeline is structured into four core analytical parts: (i) genome assembly and quality control, (ii) taxonomic classification and phylogenetic analysis, (iii) genome annotation, and (iv) advanced functional profiling. Quality control begins with read correction and trimming (SC1), followed by an adaptive genome assembly strategy integrating Canu, SPAdes, Flye, and LJA (SC2–SC5). The workflow dynamically refines genome reconstruction using an iterative assembly refinement process (SC6-SC11). Taxonomic classification (SC14–SC15) enables precise isolate identification, while genome annotation (SC16–SC18) harmonizes multi-tool gene predictions, enzyme commission (EC) numbers, COG assignments, and KEGG orthologues using hierarchical scoring systems. Functional profiling (SC19–SC27) detects plasmids, antibiotic resistance genes, CAZymes, CRISPR elements, pathogenicity islands, and insertion sequences. Each module operates within an isolated Private Process Directory (PPD) for structured data management, ensuring reproducibility through optional deterministic execution and cryptographic integrity verification (SHA-256).

Conceptually, the modules are grouped into four core analytical stages reflecting the typical progression of microbial genome analysis: i) genome assembly and quality control (SC1–SC13); ii) taxonomic classification and phylogenetic placement (SC14–SC15); iii) genome annotation (SC16–SC18), and iv) advanced functional profiling, including mobile genetic elements, antimicrobial resistance determinants, and metabolic potential (SC19–SC27). This modular architecture allows selective re-execution of individual components, while an optional deterministic mode with controlled entropy sources ensures bit-identical results across computational environments. Each processing step is encapsulated within a version-controlled container and tracked via cryptographic hashing, ensuring analytical integrity and traceability across large-scale studies.

Comprehensive microbial genome analysis begins with input-adaptive read correction and trimming (SC1) using Canu^8^. Genome assembly employs an adaptive multi-assembler strategy (SC2–SC5), integrating Canu^8^, SPAdes^9^, Flye^10^, and La Jolla Assembler (LJA)^11^. Next, assemblies are merged (MAC2^12^, SC7), circularized (Circlator^13^, SC9), and polished (Medaka/Arrow, SC11) with iterative quality assessment (QUAST^14^) and assembly selection (SC6, SC8, SC10), iteratively refining contiguity and structural accuracy. The final assembly undergoes read mapping-based coverage assessment (NGMLR^15^ and BBMap^16^, SC12) and is checked for completeness and contamination by CheckM2^17^ (SC13).

Taxonomic classification and evolutionary contextualization follow in SC14-SC15. Genomes are classified within a phylogenetic framework using GTDB-Tk^18^ (SC14). Phylogenetic trees can be constructed *de novo* to place newly assembled genomes, while also enabling the seamless integration of additional external assemblies, facilitating the generation of tailored, high-resolution phylogenetic frameworks, as demonstrated through this study (GTDB-Tk^18^, SC15, Figure 1). High-confidence genome annotation (SC16–SC18) is achieved through Bakta^18^, Prokka^20^, and MicrobeAnnotator^21^, to maximize gene prediction and functional assignment. By combining annotations from multiple tools, *StrainCascade* mitigates the idiosyncrasies of any single method, yielding more complete results than any individual tool alone. Predicted genes are unified into a non-redundant set with assigned functions (e.g. gene names, enzyme codes, COG categories, KEGG orthologs), providing a rich foundation for downstream analysis.

Advanced structural and functional profiling (SC19–SC27) reveals diverse genetic determinants of microbial behavior, ecology, and clinical significance. *StrainCascade* identifies elements of potential horizontal gene transfer, such as extrachromosomal elements and genomic islands, using PlasmidFinder^22^ and IslandPath-DIMOB^23^ (SC19, SC23). Clinical relevance is assessed via antimicrobial resistance gene detection by the ResFinder^24^ and AMRFinderPlus^25^ pipelines (SC20-SC21), while bacterial defense systems and mobile genetic elements are detected by CRISPRCasFinder^26^, ISEScan^5^, VirSorter2^27^, and DeepVirFinder^28^ (SC24-SC27). Metabolic capabilities are then profiled via KEGG metabolic modules using MicrobeAnnotator^21^ (SC18) and carbohydrate-active enzymes (CAZyme) using dbCAN3 (SC22), revealing niche occupation and adaptation strategies. The workflow concludes with the generation of an interactive HTML report (SC28, Video S1), integrating structural, taxonomic and functional genomic analyses, while also providing a comprehensive .RData file for downstream analysis and advanced exploration (Video S2). To facilitate large-scale analyses, *StrainCascade* runs each module in an isolated working directory (a ‘Private Process Directory’) that manages input/output files and prevents interference between parallel pipeline runs (Figure S1). This framework allows specific modules to be re-run independently without repeating the entire workflow, enabling efficient iterative analyses on big datasets. Together with an optional deterministic mode ensuring bit-identical outputs and cryptographic hashing for integrity verification, these features ensure *StrainCascade* provides a robust and reproducible solution for automated high-throughput bacterial genome analysis. Its modular, containerized execution and built-in safeguards enable both novice and expert users to generate high-quality assemblies and rich functional profiles across diverse datasets, establishing a strong foundation for downstream comparative genomics, evolutionary analysis, and microbial trait discovery.

### Benchmarking genome assembly performance and resource usage

To evaluate the effectiveness of genome assembly, we applied *StrainCascade* to a diverse dataset of 152 bacterial isolates (both in-house and public genomes) sequenced with PacBio RS II/Sequel II/IIe/Revio and ONT MinION/GridION platforms) (Figures 1 and 3; Table S1). We benchmarked assembly quality by comparing the results of the pipeline to assemblies produced by conventional single-assembler workflows. *StrainCascade* consistently outperformed these approaches, achieving markedly higher contiguity and more complete assemblies. In aggregate, our pipeline produced assemblies with significantly fewer total contigs per genome (adjusted p < 0.001 [q]) while maintaining greater contiguity (N50-to-genome size ratio, q < 0.001) than the alternative assemblers (Figure 3A and 3B and Table S2). These improvements indicate a higher assembly quality and a closer approach to fully closed genomes.

**Figure 3.**
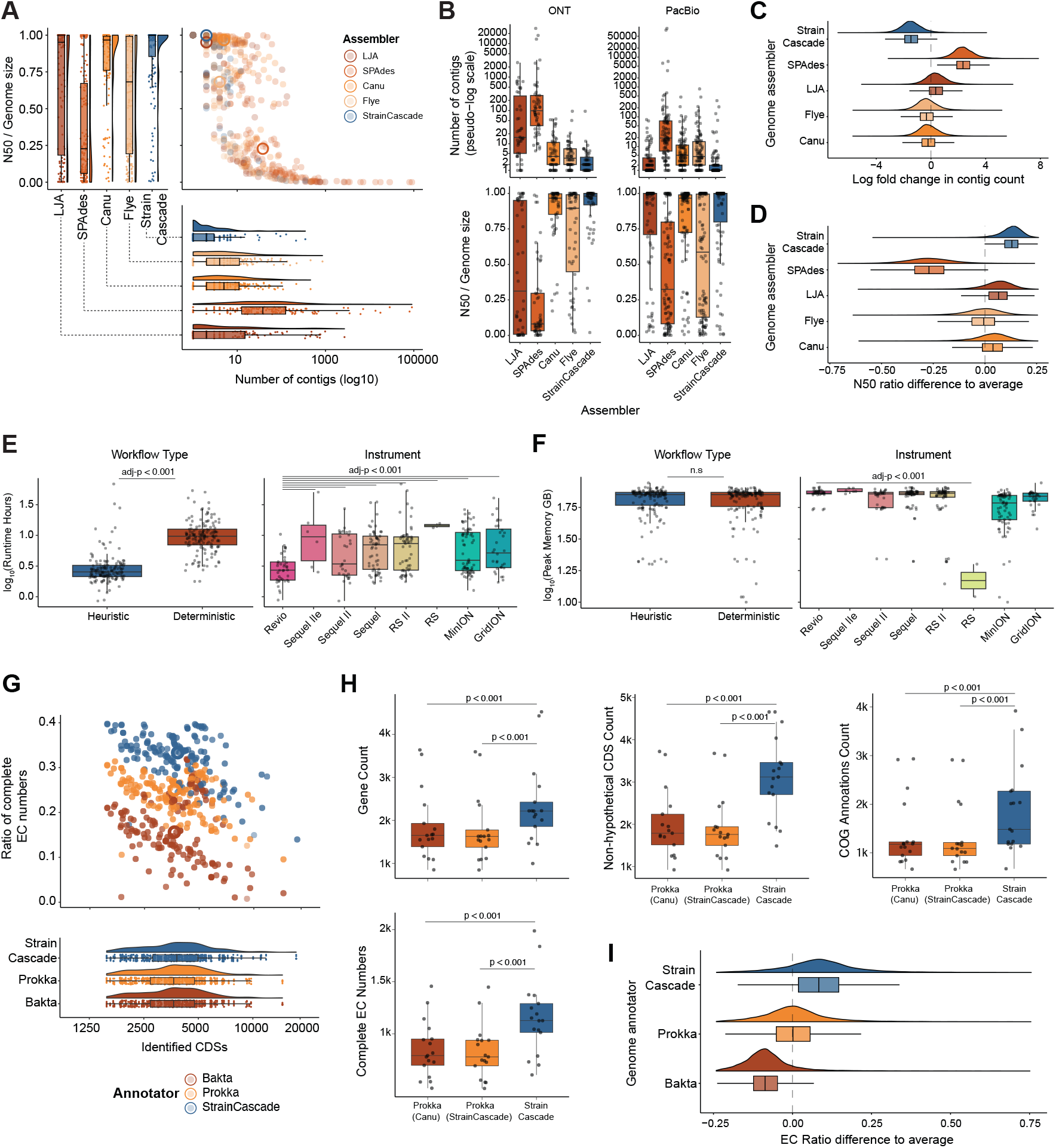
Benchmarking *StrainCascade* reveals superior genome assembly, annotation completeness, and computational efficiency compared to conventional workflows. (A) Assembly performance across different assemblers, comparing the contig count and N50-to-genome size ratio obtained using LJA, SPAdes, Canu, Flye, and *StrainCascade* (p < 0.001). (B) Assembly performance across ONT and PacBio sequencing platforms, comparing the N50-to-genome size ratio and contig count obtained using LJA, SPAdes, Canu, Flye, and *StrainCascade* (p < 0.001). (C) Posterior distribution of contig count effects, showing the log-fold change relative to the population mean. *StrainCascade* demonstrated the lowest contig fragmentation (mean = -1.43, 95% CI: -3.00 to 0.33), outperforming Canu, Flye, LJA, and SPAdes. (D) Posterior distribution of N50 ratio effects, highlighting relative improvements in contiguity across assemblers. *StrainCascade* maintained leading performance (mean = 0.12, 95% CI: -0.02 to 0.21). (E) Runtime comparison between heuristic and deterministic execution modes of *StrainCascade* across platforms. Deterministic runs incurred longer runtimes (p < 0.001). (E) Peak memory usage across heuristic and deterministic runs, stratified by sequencing instrument. Memory usage remained modest and predictable across platforms, with no major difference between modes. (G) Number of CDSs and complete EC number assignments across all 152 genomes, comparing Bakta, Prokka, and *StrainCascade* annotations. (H) Comparative annotation metrics for in-house isolates sequenced with PacBio Revio, showing that *StrainCascade* predicted more CDSs, more complete EC numbers, and more COG annotations per genome than Prokka alone (p < 0.001). (I) Posterior distribution of effects on complete enzyme commission (EC) number ratios, illustrating annotation completeness across genome annotators. The *StrainCascade* pipeline provides the highest EC annotation recovery (mean = 0.09, 95% CI: -0.15 to 0.37) compared to the population mean, outperforming Bakta and Prokka. Comparison of genome assembly metrics across sequencing platforms (PacBio Revio, Sequel, Sequel II, RS II; ONT GridION, MinION) and assemblers (LJA, SPAdes, Canu, Flye, *StrainCascade*). Error bars represent the standard deviation of 95% credible intervals, as appropriate. Statistical significance was assessed using hierarchical Bayesian modeling or Wilcoxon signed-rank tests with FDR correction.

Indeed, hierarchical Bayesian modelling showed that *StrainCascade* achieved the lowest contig counts in all posterior draws (P(best) = 1.00, reflecting its multi-stage optimization architecture that systematically selects and refines the most promising assemblies. Effect size estimation by (conservative) Bayesian modelling indicated a trend for reduced contig numbers (estimated a log-fold change of -1.43; 95% credibility intervals [CI]: -3.00 to 0.33) (Figure 3C and Table S3). Superior assembly contiguity was confirmed as well: *StrainCascade* showed consistently superior posterior draws (P(best) = 1.00) and increased the N50-to-genome-size ratio by ∼0.15 (95% CI: –0.02 to 0.21) (Figure 3D), reflecting longer continuous assembled sequences relative to total genome size. Notably, these gains were observed consistently across both PacBio and ONT datasets, making *StrainCascade* the top performer in assembly quality among all tested pipelines, delivering the most contiguous assemblies with the fewest scaffolds (Figure 3A-D and Figure S2A-2C).

To assess computational performance, we evaluated runtime and memory usage using linear mixed-effects models. Two execution modes were benchmarked: heuristic, which prioritizes speed with flexible parallelization, and deterministic, which enforces strict reproducibility by fixing random seeds, standardizing entropy sources, and running in single-threaded mode. *StrainCascade* runtime varied significantly by workflow type, with deterministic runs taking longer on average than heuristic runs (Figure 3E). Instrument platform was also a significant predictor, with assemblies from ONT (e.g., MinION, GridION) generally requiring more time than PacBio Revio (e.g., Sequel, RS II; all *q* < 0.001) (Figure 3E). These results suggest that both pipeline settings and sequencing technology influence computational runtime. In contrast, genome size and input file size had minimal explanatory power, suggesting runtime is more sensitive to platform complexity and deterministic processing constraints than to data volume (Figure S2D).

Peak memory usage was modest overall (typically between 57 GB and 74 GB) and slightly lower for ONT-based datasets compared to PacBio platforms (e.g., Revio, Sequel II/IIe) (Figure 3F). This likely reflects differences in read length and error-correction complexity between the two technologies. Workflow type had no significant effect on memory usage (Figure 3F), and neither input file size nor genome size explained substantial variance (Figure S2E), indicating that memory requirements remain modest and consistent across data types. These findings underscore the trade-off between reproducibility and runtime while showing that *StrainCascade* operates with modest and consistent resource demands across platforms.

### Enhanced genome annotation accuracy

*StrainCascade* enhanced genome annotation compared to existing tools. Within the 152 bacterial isolates, the pipeline consistently identified more coding sequences (CDS) and assigned Enzyme Commission (EC) numbers than either Bakta or Prokka alone (Figure 3G). When benchmarked specifically for in-house isolates sequenced on PacBio Revio (Table S1), *StrainCascade* predicted a higher number of CDS per genome and achieved a greater fraction of genes with assigned EC numbers and COG (Clusters of Orthologous Groups) annotations compared to annotations generated by Prokka (Figure 3H). We observed superior EC number completeness, with a relative improvement of 0.09 (95% CI: -0.15 to 0.37) and P(best) = 1.00 in an additional Bayesian hierarchical regression. These results underscore the scalability and accuracy of the pipeline, establishing *StrainCascade* as an optimal solution for large-scale microbial genomics (Figure 3I and Table S2). By combining the outputs of multiple annotation engines, the pipeline leverages complementary databases and prediction algorithms, yielding a more comprehensive gene catalogue for each genome.

### Resolving complex genomes and unclassified isolates

Beyond aggregate performance metrics, we next probed the capacity of the pipeline to resolve taxonomically ambiguous or structurally complex bacterial genomes, those that are typically unclassifiable or refractory to standard reconstruction approaches. For example, *Veillonella nakazawae*, an isolate from our lab unidentifiable by MALDI-TOF, was classified through high-contiguity assembly and phylogenetic placement (Figure 4A). In another case, *Vibrio cholerae* was fully assembled into two circular chromosomes, allowing comprehensive gene annotation, antimicrobial resistance (AMR) profiling, and mobile element detection (Figure 4A). These examples illustrate how the pipeline can reconstruct complex genomes and resolve previously unclassifiable isolates.

**Figure 4.**
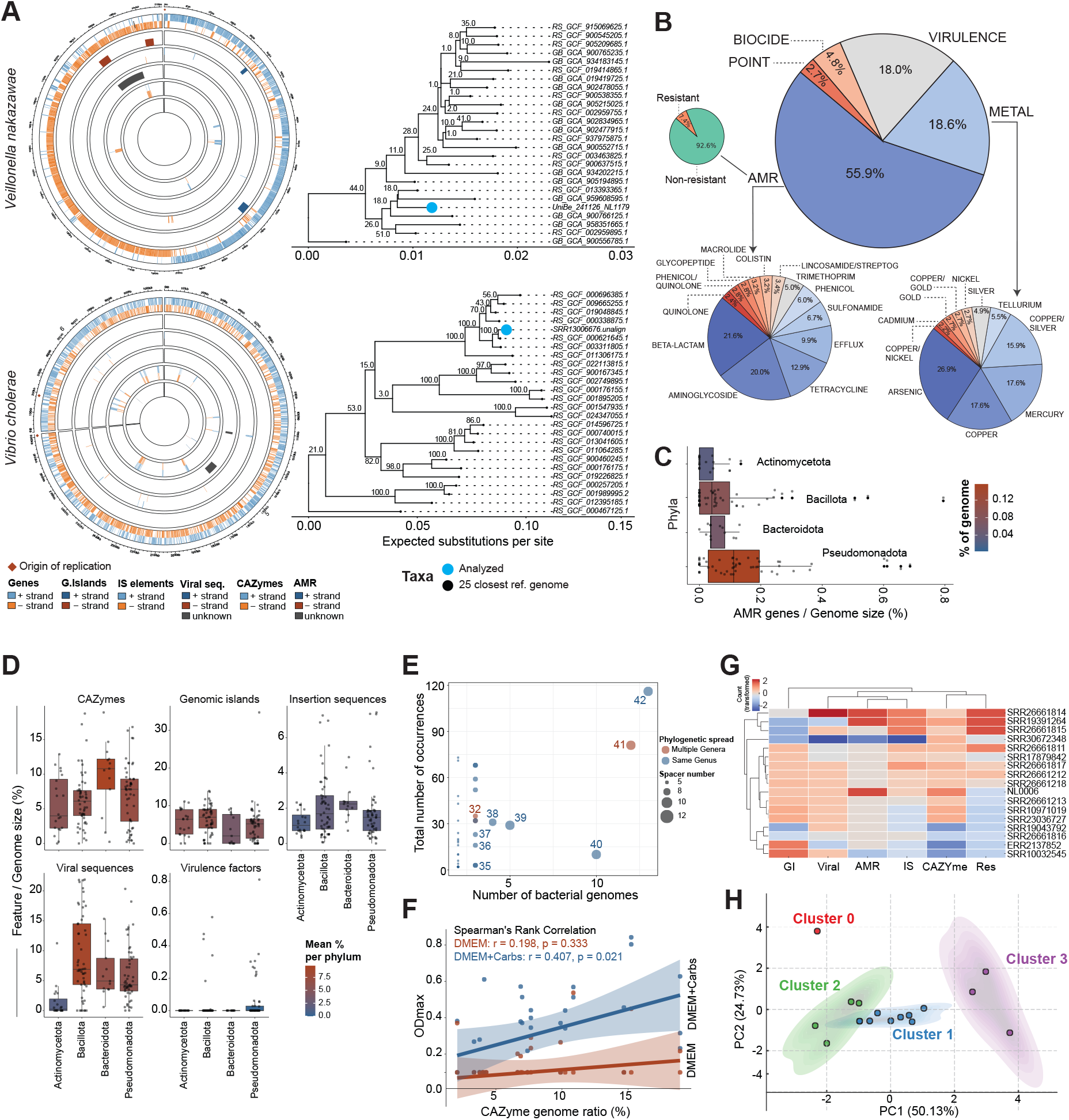
*StrainCascade* enables strain-resolved genomic reconstruction, functional trait mapping, and discovery of adaptive signatures across microbial populations. (A) Phylogenetic resolution and genome reconstruction, showing how *StrainCascade* identifies and assembles novel isolates, including *Veillonella nakazawae* (unclassified by MALDI-TOF) and *Vibrio cholerae* with its two circular chromosomes. (B) Functional resistance profiling summarizing the distribution of AMR genes, metal resistance, virulence factors, biocide resistance, and point mutations across all analyzed bacterial genomes. The large pie chart represents the mean (%) distribution of these functional resistance categories. Below the large pie chart, the left pie chart illustrates the percentage distribution of antimicrobial resistance classes, while the right pie chart depicts the percentage distribution of metal stress resistance classes. Additionally, a smaller pie chart on the left side of the large pie chart displays resistance and non-resistance profiles identified using ResFinder, which detects acquired AMR genes and chromosomal mutations. (C) Taxonomic association of AMR genes, demonstrating that Pseudomonadota harbors a nominally higher proportion of AMR genes relative to genome size among the four most abundant phyla. (D) Comparative genomic analysis across phyla, highlighting the distribution of CAZymes, genomic islands, insertion sequences, viral sequences and virulence factors. To examine the influence of bacterial phylum on feature-specific ratios, a linear model (lm) was fitted with phylum (Pseudomonadota as reference) as a fixed effect, while Kruskal-Wallis tests assessed global differences, with p-values adjusted using the false discovery rate (FDR) method. (E) CRISPR spacer distribution across bacterial genomes, illustrating the frequency of shared vs. unique spacers. Most spacers remain genus-specific, with a small subset shared across species, suggesting lineage-specific immune memory against phages. (F) Correlation plot of the growth of 17 isolates (NL0006, NL0022, NL0034, NL0042, NL0051, NL0064, NL0092, NL0123, NL0130, NL0141, NL0165, NL0267, NL0381, NL0399, NL0561, NL1066, NL1179) with different CAZyme gene profiles when cultured in DMEM and DMEM + carbohydrates. (G) Heatmap showing strain-specific genomic plasticity in *E. coli* with the distribution of AMR genes, genomic islands, insertion sequences, ResFinder-resistant classes (Res), viral elements, and CAZymes. (H) Unsupervised K-means clustering (K=4) *of E. coli* strains based on genomic features, identifying four functional groups on a PCA plot.

### Lineage-level diversity in genome plasticity and metabolic potential

We next tested the capacity of the pipeline to reveal lineage-specific trends in genome architecture and functional potential across 152 diverse bacterial strains. Because the pipeline simultaneously generates assemblies, annotations, and functional profiles for all genomes, it enables high-resolution comparative analyses that reveal how genomic traits vary across taxa. *StrainCascade* identified a wide array of AMR genes as well as virulence factors, metal resistance genes, and biocides in 152 analyzed strains (Figure 4B). We demonstrate variation in the ratios of the genome dedicated to AMR between phyla (Figure 4C) and lineage-specific diversity in AMR genes, CAZymes, insertion sequences, and genomic islands across the dataset, reflecting the unique resistance potential of each taxon (Figure 4D and Figure S3A-D and S4). For example, the phylum Bacillota had the highest content of viral sequences, whereas Actinomycetota exhibited significantly lower levels (p = 0.002). Bacteroidota genomes were enriched in CAZyme genes (p = 0.039), and Pseudomonadota contained the most virulence factors (p = 0.005) (Figure 4D). Pronounced strain-level variation was also observed: *Klebsiella pneumoniae, Escherichia coli, Salmonella enterica*, and *Staphylococcus aureus each* showed distinct profiles of AMR genes, CAZyme repertoires, and mobile genetic elements (plasmids, insertion sequences, genomic islands), highlighting the considerable genomic variability in strains from same bacterial species (Figure S3A-D).

*StrainCascade* also profiles lesser-explored facets of genome diversity such as CRISPR-Cas spacer content. CRISPR spacers can be thought of as bacterial “mugshots” of past viral encounters – each spacer is a sequence derived from a virus (phage) that infected the bacterium or its ancestors, providing immunity against future infection by the same phage. Across the 152 genomes, *StrainCascade* identified a total of 344 unique CRISPR spacers. Out of 344 unique spacer sequences, 42 occurred in >1 genome, suggesting that a few bacteriophages exert broad selective pressures across multiple hosts. Our analysis of recurrent spacer sequences revealed near-exclusive genus or species-specific patterns for most CRISPR spacers and only two spacers were shared across genera (Figures 4E and S3E). Consistent with sporadic viral exposure, most spacers appeared only once or twice per genome; however, a handful recurred up to 94 times, indicating repeated encounters with the same virus (Figure 4E).

Together, the comparative analyses above establish that *StrainCascade* can capture lineage-specific functional and structural variation at multiple levels, from broad phylum trends down to fine-scale differences between individual strains, thereby revealing the genomic signatures of adaptation, ecology, and evolutionary history across our collection of bacteria.

### Linking CAZyme profiles to metabolic phenotypes

Recognizing that genome content underpins metabolic capability, one can use *StrainCascade* to link genomic architecture with functional traits, paving the way for systematic studies of microbial adaptation. To probe the functional impact of carbohydrate-active enzymes (CAZymes), each isolate was cultured in a defined minimal medium (DMEM) with or without supplemental complex carbohydrates, and growth was monitored to assess metabolic response (Figures 4F and S5). Strains annotated by *StrainCascade* as CAZyme-rich exhibited markedly enhanced growth in carbohydrate-supplemented media, while those with limited CAZyme repertoires showed minimal to no growth improvement (Figures 4F and S5). This illustrates that possessing an expanded set of carbohydrate-active enzymes confers a metabolic advantage when polysaccharides are available. This finding validates that the genomic predictions made by *StrainCascade* have tangible phenotypic consequences.

### Strain-level functional clustering reveals adaptive signatures

Phylogenetic proximity does not always reflect functional similarity, especially in genomes shaped by horizontal gene transfer and niche-specific adaptation. To identify patterns of functional adaptations beyond phylogenetic relationships, we employed unsupervised clustering of functional genetic markers characterized by *StrainCascade* to reveal strain-specific evolutionary trajectories. We selected a set of 17 Escherichia coli genomes from our collection as a test case and compiled a matrix of their genomic features of interest: total counts of unique AMR genes, CAZyme families, insertion sequences, genomic islands, and other mobile or adaptive elements characterized by the pipeline (Figures 4G, 4H and S6). This *E. coli* set exhibited extensive genetic variability. In total, the 17 strains collectively harbored 312 unique AMR genes (1029 total occurrences) and genes spanning four major CAZyme families (6915 entries). These genomes also contained 3306 distinct insertion sequences, 1359 genomic islands, and 98 unique resistance determinants (15288 occurrences) (Tables S4-S9). Unsupervised clustering (k=4, Figure S6E) revealed four functionally distinct groups (Figures 4H and Table S10). Cluster 0 (‘stability adaptors’) had moderate AMR gene content and few genomic islands, indicating a balance between genomic stability and resistance acquisition. Cluster 1 (‘mobile element enriched’) was enriched in insertion sequences and genomic islands, suggesting frequent horizontal gene transfer. Cluster 2 (‘metabolic innovators’) displayed the highest genomic island content alongside elevated CAZyme levels, suggesting metabolic adaptations by both horizontal gene transfer-driven and intrinsic genetic diversification. Cluster 3 (‘resistance reservoirs’) harbored the highest AMR gene burden and the most diverse resistance classes, representing high-risk, clinically relevant strains. These functional clusters, derived purely from genomic content profiles, are notable because they do not strictly correspond to the phylogenetic relationships of the *E. coli* strains. In other words, strains that are distantly related in evolutionary terms are sometimes clustered together functionally, whereas closely related strains fall into different functional clusters. This suggests that horizontal gene transfer and selective pressures have reshaped certain genomes in parallel, leading to the convergent acquisition of traits. The integrated analysis of the pipeline can make it straightforward to generate the high-dimensional data needed to uncover adaptive patterns invisible to traditional taxonomy, generating testable hypotheses about strain-specific evolution and ecological adaptation.

## DISCUSSION

*StrainCascade* integrates high-throughput reconstruction and characterization of bacterial genomes from long-read sequencing data, delivering superior assembly quality, comprehensive functional annotation, and comparative phylogenetic analysis, thus enabling precise strain-level investigations. Its modular, deterministic design with adaptive module selection guarantees reproducible and robust genome reconstruction across diverse sequencing platforms (Figure 2 and 3). *StrainCascade* advances microbial surveillance, epidemiology, and evolutionary genomics by uncovering strain-specific adaptations and genomic plasticity crucial for understanding host-microbe interactions.

Beyond performance benchmarking, our results demonstrate how *StrainCascade* enables functional and evolutionary insight at multiple levels of resolution, from species-wide annotation improvements to strain-specific differences in mobile elements and metabolic potential (Figure 4). The ability to resolve complex structural features, assign high-confidence functions, and reveal latent genomic traits such as resistance potential or niche adaptation is particularly valuable in the context of public health, microbial ecology, and host-microbiome research. By providing more complete assemblies and richer annotations, *StrainCascade* allows researchers to ask deeper questions about each genome. For example, understanding that a given strain has an expanded arsenal of AMR genes or unique metabolic pathways can inform hypotheses about its ecological niche or clinical risk, which would be missed if the genome were fragmented or poorly annotated.

Importantly, the increase in annotated genes observed with *StrainCascade* does not reflect inflated predictions but rather a more complete and functionally meaningful recovery of true coding sequences. Individual tools differ in stringency and database scope; for instance, Bakta tends to call slightly more genes than Prokka by default but may fragment open reading frames or assign more hypothetical labels^19^. By integrating annotations from Bakta and Prokka and supplementing with pathway-level insights from MicrobeAnnotator^21^, *StrainCascade* captures complementary strengths while mitigating tool-specific biases. This approach mirrors recent efforts like the Beav pipeline^29^, which showed that augmenting Bakta with additional modules significantly reduced hypothetical annotations. Similarly, the unified annotation strategy of *StrainCascade* recovers functionally annotated genes that might otherwise be missed or labeled as uncharacterized, thereby improving the interpretability of microbial genomes.

The superior assembly outcomes of *StrainCascade* stem from its multi-assembler consensus strategy (Figure 2) and rigorous refinement steps. By merging results from multiple long-read assemblers, the pipeline captures a more complete and accurate genome representation than any single-tool approach. This echoes the benefits of manual consensus tools like Trycycler^30^, which improves assembly accuracy by combining multiple drafts but unlike Trycycler, *StrainCascade* achieves these gains in a fully automated, reproducible, and deterministic manner. Circularization and refinement modules further enhance quality, frequently producing closed, publication-grade genomes without manual intervention. For example, *Vibrio cholerae* was reconstructed into two circular chromosomes in a single pass. This is particularly impactful given that even recent long-read assemblers often miss small plasmids or other structurally complex features. By combining complementary tools and rigorous post-assembly processing, *StrainCascade* maximizes contiguity and completeness, improving downstream analyses such as variant detection, comparative genomics, and functional annotation.

The modular architecture of *StrainCascade* offers fertile ground for future expansion (Figure 2). Upcoming modules could incorporate host-interaction modeling, metagenomic binning, pan-genome reconstruction for large strain collections, or integration with epigenomic data (to include methylation patterns from long reads, for instance). The containerized execution model ensures that such extensions can be developed and deployed without disrupting core functionality, creating a sustainable path toward more comprehensive multi-omic microbial analyses.

Furthermore, *StrainCascade* is well-positioned to support applications in synthetic biology and microbial therapeutics. The ability to precisely reconstruct and functionally annotate bacterial genomes enables the rational engineering of strains with tailored properties, whether for microbiome modulation, probiotic design, or synthetic community construction. In combination with host-microbe modeling, *StrainCascade* lays the groundwork for next-generation therapeutic development grounded in high-resolution genomic data. Its integration of CRISPR-Cas analysis and horizontal gene transfer detection further provides a robust framework for biosafety profiling and strain optimization, facilitating the design and monitoring of clinically relevant microbes in translational and regulatory settings (Figure 4).

Together, these improvements make *StrainCascade* a versatile, user-friendly, functionally rich, and reproducible solution for high-throughput, long-read bacterial genome analysis. By enabling precise genome reconstruction, deterministic execution, and modular expansion, *StrainCascade* provides a sustainable platform for both exploratory and translational applications in microbiology. *StrainCascade* is fully open-source and modular by design, supporting community-driven extensions and transparent benchmarking. As bacterial genomics enters an era of increasing scale and functional complexity, such collaborative frameworks will be essential for ensuring analytical continuity, reproducibility, and shared progress across research groups and clinical consortia.

### Limitations of the study

While *StrainCascade* provides a comprehensive and reproducible framework for long-read bacterial genome analysis, several limitations warrant consideration. The current implementation is optimized for isolated genomes sequenced using PacBio and Oxford Nanopore technologies, with support for Illumina-based short-read data limited to post-assembly inputs. Given the availability of robust, dedicated pipelines for short-read analysis^3-6^, we did not extend *StrainCascade* to natively handle short-read workflows. Second, although the pipeline is platform-agnostic, its performance on lower-depth or highly fragmented datasets remains to be systematically benchmarked. Users working with environmental or host-associated samples where DNA quality or yield is limited may require additional pre-processing or error correction not yet integrated into *StrainCascade*. Third, although *StrainCascade* enhances functional annotation, its profiling capabilities remain dependent on reference databases (e.g., KEGG, CAZy, ResFinder), which are inherently biased toward well-characterized organisms. This may limit the interpretability of novel genes or pathways in poorly characterized taxa, particularly in environmental or extremophile datasets. Future development could incorporate machine-learning-based functional prediction and structure-informed annotation to expand coverage beyond current database constraints. Finally, while the deterministic mode ensures reproducibility, in most cases, it increases runtime substantially compared to heuristic execution. This trade-off may be a limiting factor in real-time surveillance or large-scale screening scenarios unless computational resources are abundant, or optimized batch strategies are adopted.

## RESOURCE AVAILABILITY

### Lead contact

Further information and requests for resources should be directed to and will be fulfilled by the lead contact, Bahtiyar Yilmaz.

### Materials availability

This study did not generate new unique reagents.

## DATA AND CODE AVAILABILITY

All sequencing data from the public NCBI SRA are accessible via accession numbers listed in Table S1 (Sample IDs). All newly generated samples from our laboratory (Sample IDs in the format NLXXXX, starting with NL) are available for download from the Zenodo repository: https://doi.org/10.5281/zenodo.14921436 including raw reads (FASTQ), assemblies (deterministic run 2), interactive HTML reports (deterministic run 2), and run reports (deterministic run 2).

All models, codes, and notebooks to reproduce our analysis and figures are available at GitHub: https://sbujordi.github.io/StrainCascade_documentation/ and https://github.com/SBUJordi/StrainCascade. The code used to benchmark workflow efficiency and analysis has been archived and is available on Zenodo at: DOI: 10.5281/zenodo.14921436. All preprint supplementary files can be downloaded at https://zenodo.org/records/18479044.

## ACKNOWLEDGMENTS

We thank all subjects and patients for their commitment and donation of biological samples. We are grateful for the resource on High-Performance Computing Cluster UBELIX of the University of Bern. We are thankful to Dr. Nataliia Polishchuk for her constant support with microbiological applications. We are deeply grateful to the Dr. Pamela Nicholson and University of Bern NGS Platform for performing the PacBio WGS sequencing for in-house bacterial isolates. Viable bacterial strains isolated in this study are also available from the corresponding author for further studies.

## FUNDING

We thank the European Crohn’s and Colitis Organisation (ECCO) for funding this study via the ECCO Grant 2023 and the Bern Center for Precision Medicine (BCPM), University of Bern (Recipient – B.Y.). B.Y. was also supported by the Swiss National Science Foundation (SNSF) Starting Grant: TMSGI3_211300, and B.M. was supported by SNSF Project Grant: 320030_185286. S.B.U.J. was supported by the Swiss National Science Foundation (SNSF) MD-PhD grant: 323630_221867.

## AUTHOR CONTRIBUTIONS

B.Y. conceived and supervised the project together with B.M. S.B.U.J. designed and developed the pipeline. S.B.U.J. performed all analyses with the help of I.B.. J.L., and N.F. helped with bacterial isolation, culturing and growth dynamics experiments. S.B.U.J. wrote the first draft of the manuscript with B.Y. All authors wrote and contributed to the manuscript. The authors read and approved the final manuscript.

## DECLARATION OF INTERESTS

The authors declare no competing interests.

## STAR+METHODS

Detailed methods are provided in the online version of this paper and include the following:

- KEY SOURCE TABLE
- RESOURCE AVAILABILITY
  - Lead contact
  - Material availability
  - Data and code availability
- SUBJECT DETAILS
  - Study design
  - Ethics statement
- METHOD DETAILS
  - *StrainCascade* setup and execution pipeline
  - Wrapper-based execution for streamlined pipeline workflows.
  - Automated pipeline installation and environment setup.
  - Reproducibility and computational environment
  - Cross-platform dataset selection and composition
  - Taxonomic Diversity
  - In-house bacterial isolates and sequencing with PacBio Revio platform
  - Stoma sample collection and bacterial isolation
  - Identification and preservation
  - DNA extraction and WGS sequencing
  - Reads correction and trimming
  - High-precision deterministic genome assembly
  - Genome assembly
  - Assembly assessment
  - Assembly refinement
  - Assembly selection
  - Taxonomic identification and *de novo* classification
  - Classify workflow
  - *De novo* workflow
  - Plasmid identification
  - Genome annotation
  - Bakta annotation
  - Prokka annotation
  - MicrobeAnnotator annotation
  - Annotation processing
  - Functional annotation integration
  - Genomic islands identification
  - Metabolic pathway analysis
  - Antimicrobial resistance gene analysis
  - Characterization of genome plasticity
  - CRISPR-Cas systems detection
  - Insertion sequence (IS) elements
  - Viral sequence detection
- QUANTIFICATION AND STATISTICAL ANALYSIS
  - Non-parametric comparisons of assembly and annotation performance
  - Hierarchical Bayesian modeling
  - Posterior analysis and reporting
  - Computing resource

## KEY SOURCE TABLE

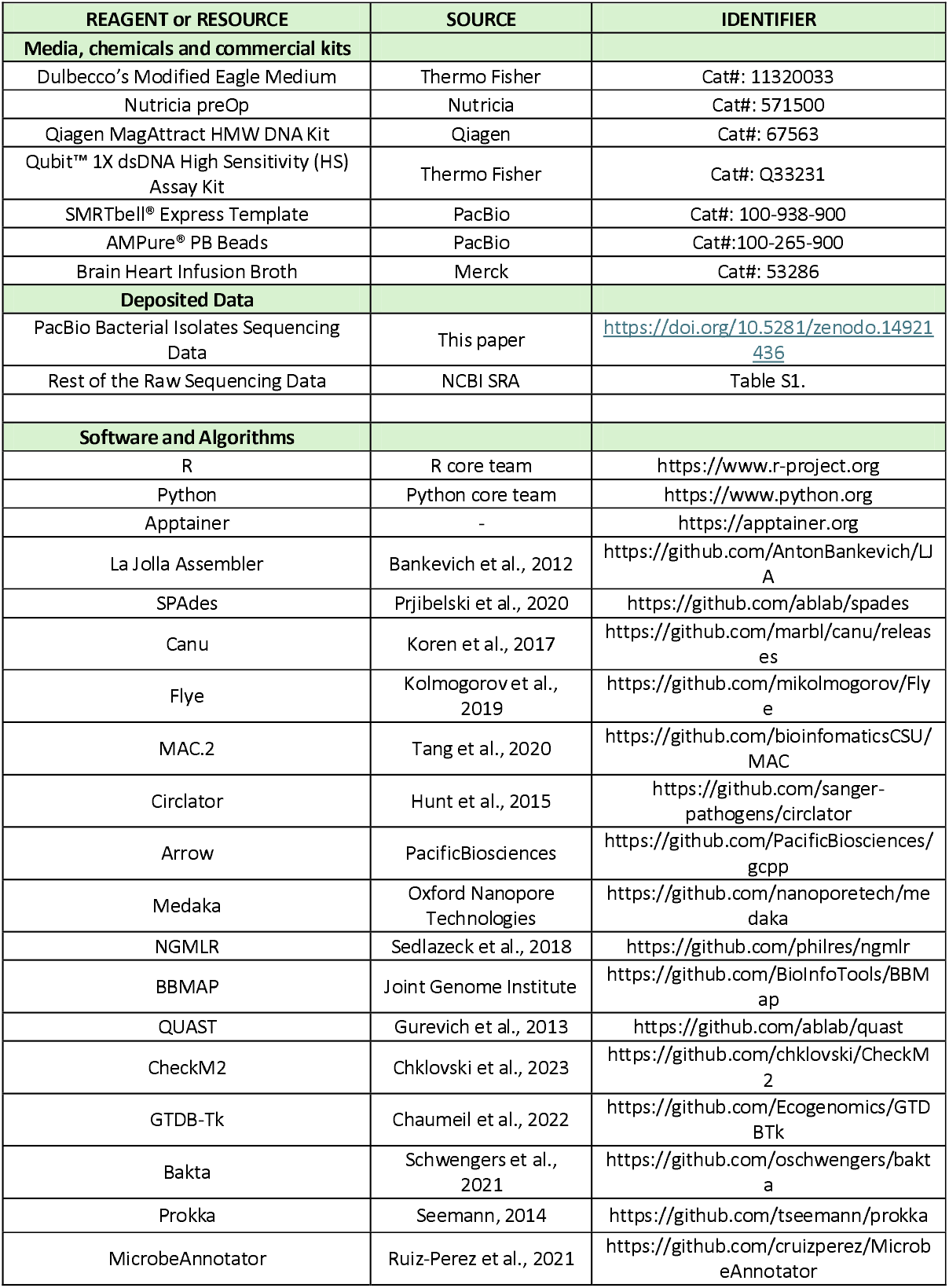
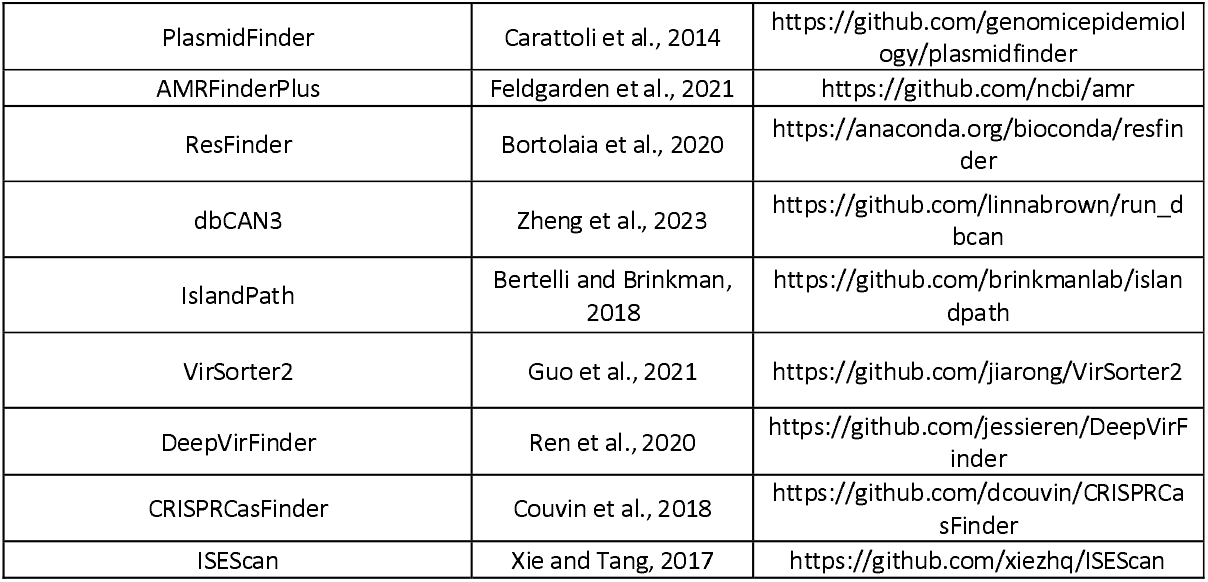

## MAIN FIGURES

## SUPPLEMENTAL FIGURES

**Figure S1.**
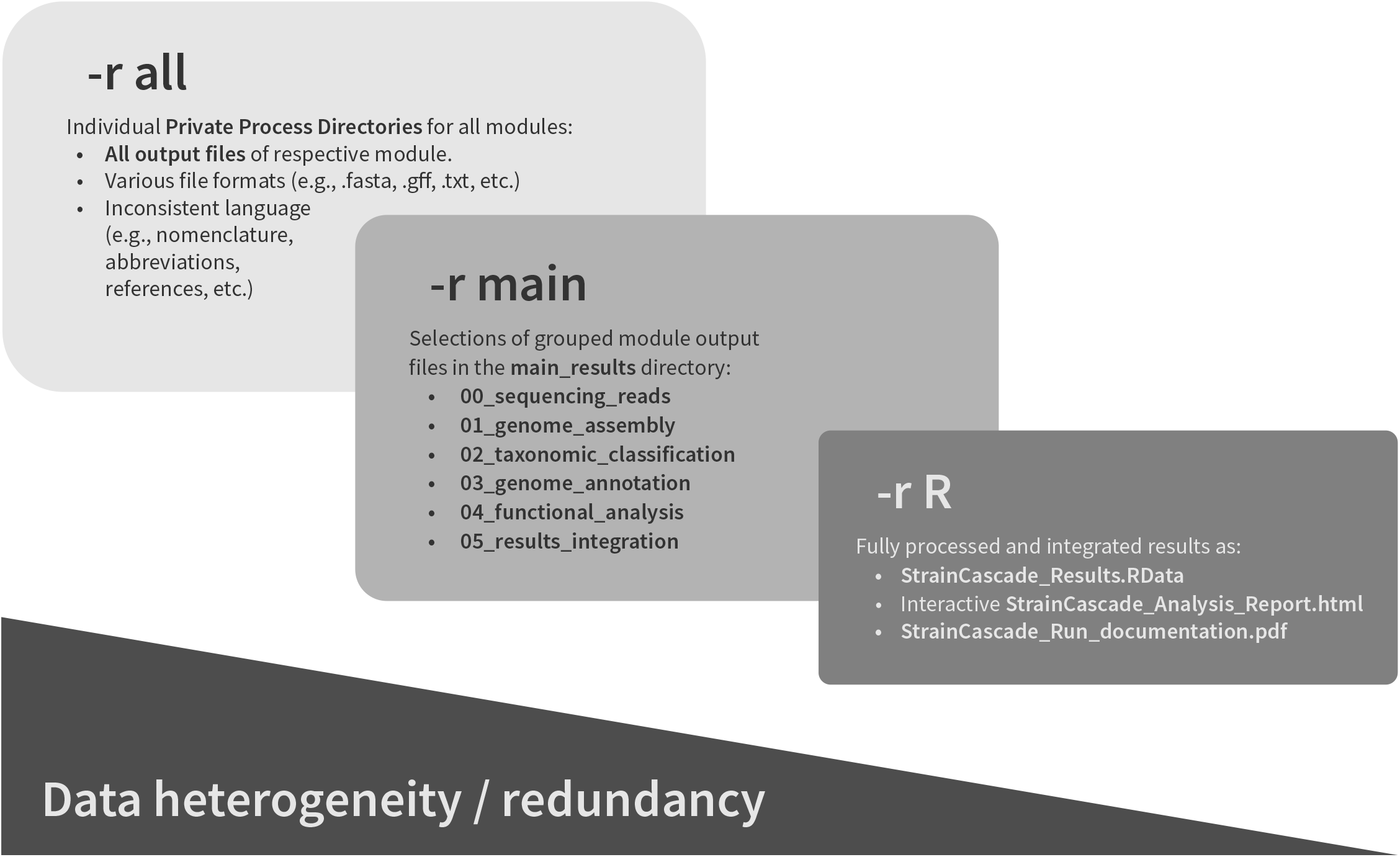
The execution framework of *StrainCascade* ensures reproducibility and structured data management. Each module operates within an isolated Private Process Directory (PPD), systematically managing intermediate and final files to maintain workflow integrity. Users can customize result retention granularity with three options: *all, main*, or R. The pipeline has the option of deterministic execution with controlled entropy sources, ensuring reproducibility across computational environments. Automated cryptographic integrity verification (SHA-256 hashing) safeguards data consistency throughout the workflow. Additionally, *StrainCascade* supports adaptive multi-threading execution, optimizing resource allocation based on system availability. This structured computational framework enables seamless scalability from single-genome assembly to large-scale comparative genomics while maintaining full documentation of pipeline execution for enhanced transparency.

**Figure S2.**
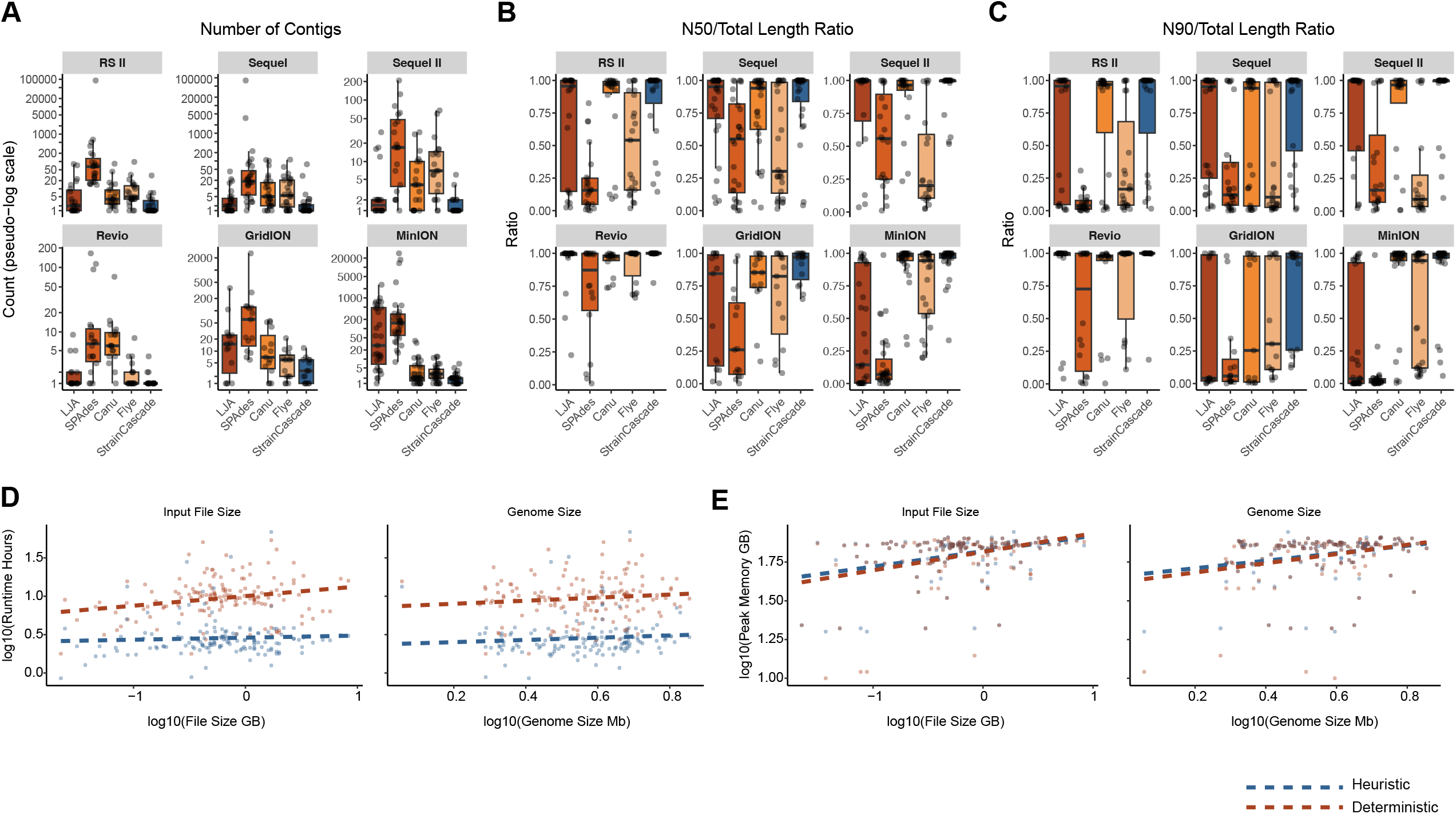
Benchmarking *StrainCascade* across assemblers, sequencing platforms, and computational performance parameters. (A) Number of contigs per assembly across different sequencing technologies and assemblers, presented on a pseudo-log scale. (B) N50-to-total genome length ratio, highlighting assembly contiguity across platforms. (C) N90-to-total genome length ratio, demonstrating further contiguity improvements. (D) Relationship between input file size (log_10_ scale) and *StrainCascade* runtime (log_10_ hours) and relationship between genome size (log_10_ Mb scale) and *StrainCascade* runtime (log_10_ hours). (E) Peak memory usage (log_10_ GB) as a function of input file size and genome size. Error bars represent standard deviation.

**Figure S3.**
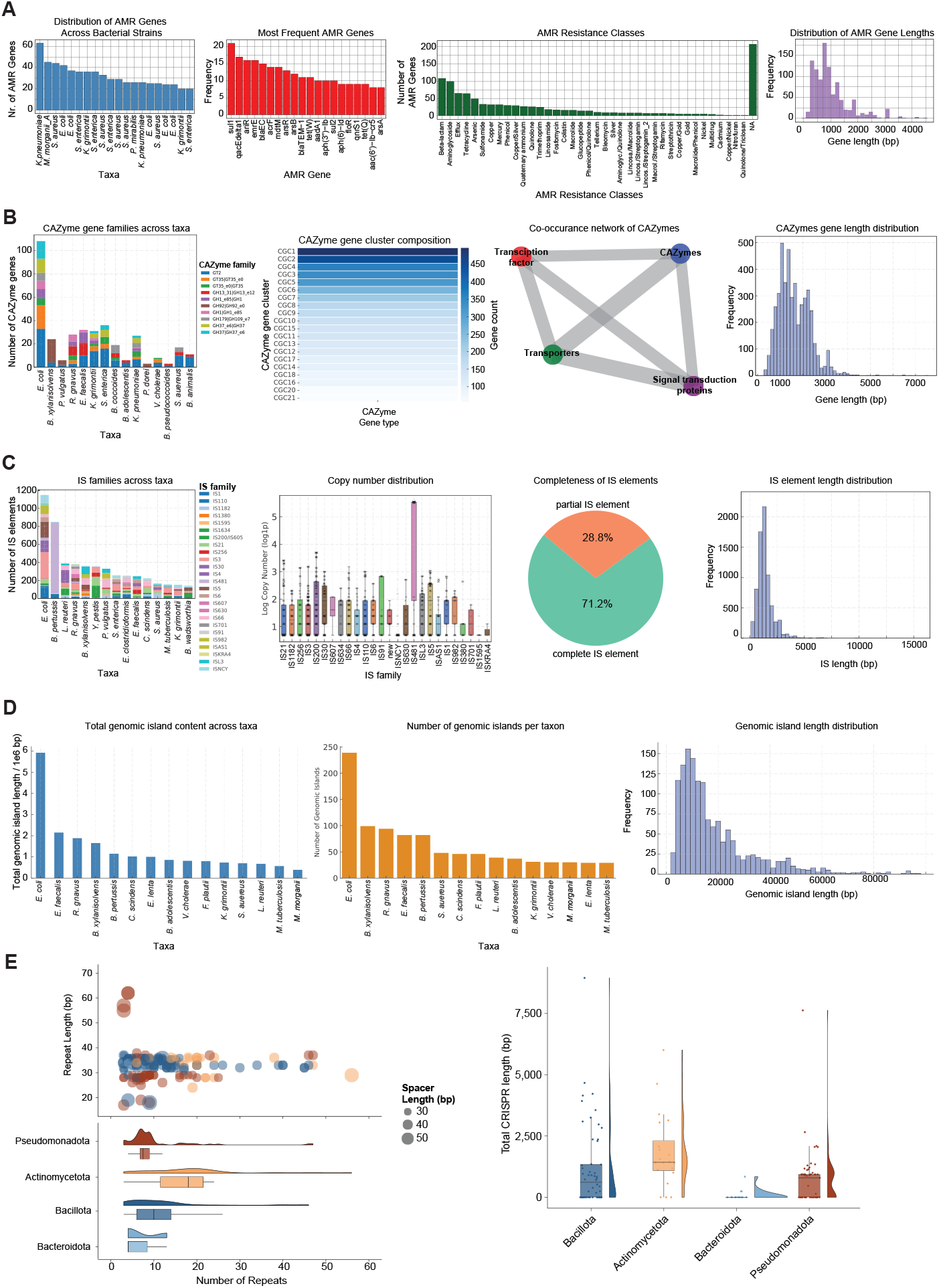
Detection of genomic functional features across bacterial strains. (A) AMR genes are detected across bacterial strains, revealing strain-specific variations in abundance, identifying the most prevalent resistance genes, classifying them into resistance classes, and analyzing their length distribution. (B) CAZyme genes are analyzed across bacterial taxa, revealing their distribution, co-occurrence with transporters, transcription factors, and signaling proteins, gene length diversity, and the composition of CAZyme gene clusters, highlighting key functional associations. (C) Insertion sequences (IS) are analyzed across bacterial taxa, examining their distribution, copy number variations, proportion of complete versus partial elements, and length diversity across strains. (D) Genomic islands are analyzed across bacterial taxa, assessing their total content per taxon, species-specific abundance, and length distribution, providing insights into genomic plasticity and adaptation. (E) CRISPR spacer distribution and cross-genome conservation are shown using the number of repeats and repeat length (bp) across major bacterial phyla, including Pseudomonadota, Bacteroidota, Actinomycetota, and Bacillota (left panels). Box-dot plot with a histogram showing the total CRISPR length (bp) distribution across these major bacterial phyla (right panel).

**Figure S4.**
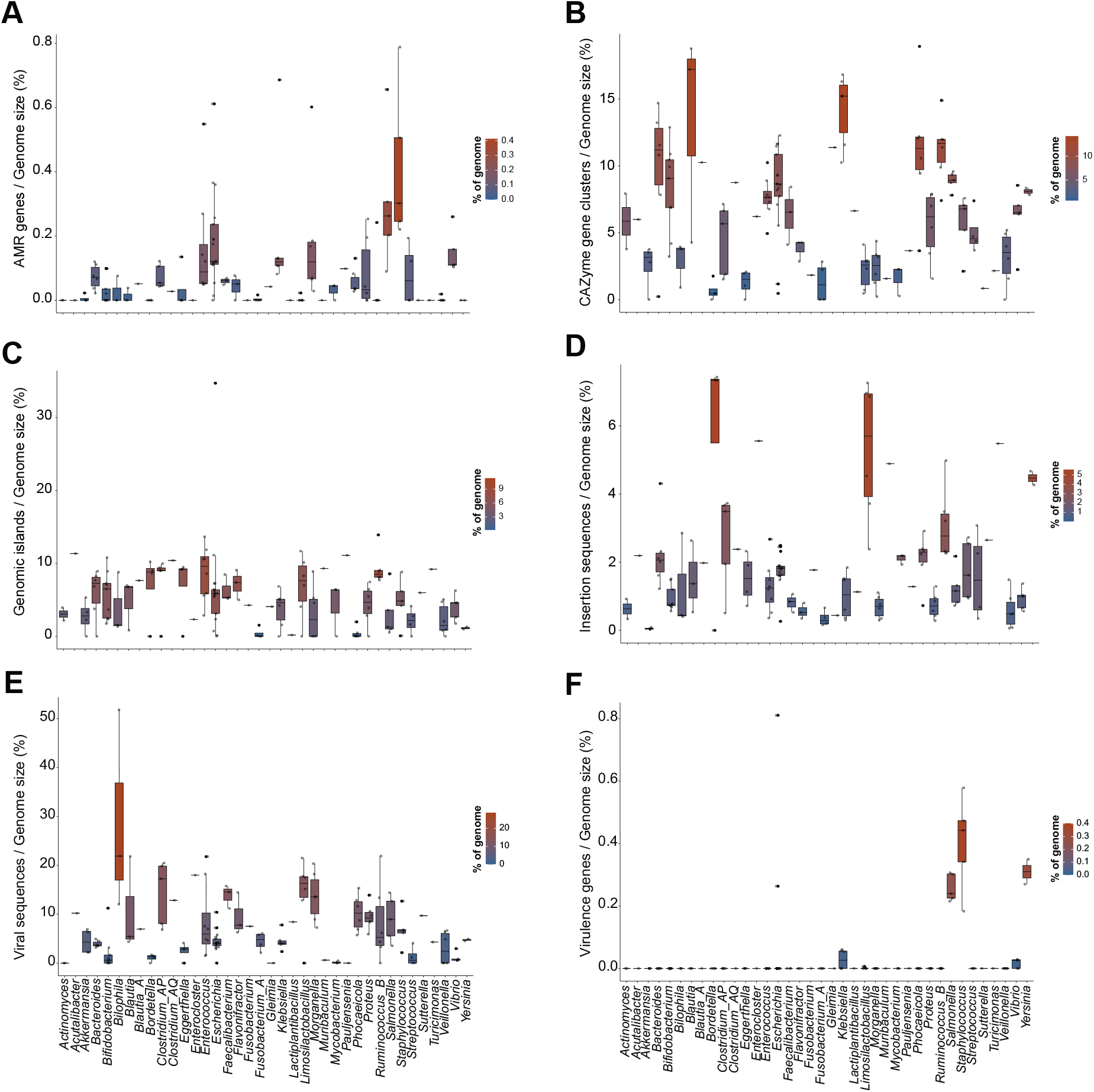
Taxa-specific variability in functional and mobile element distribution. (A) Proportion of AMR genes per genome size across bacterial taxa, highlighting variability in resistance potential among different species. (B) CAZyme gene cluster density, showing the relative abundance of carbohydrate-active enzyme clusters across genomes. (C) Genomic islands per genome size, illustrating the distribution of horizontally acquired gene regions across different taxa. (D) Insertion sequence elements per genome size, reflecting the extent of transposable elements contributing to genome plasticity. (E) Viral sequences per genome size, showing variation in the prevalence of viral insertions among bacterial genomes. (F) Virulence gene density, indicating differences in the genetic potential for pathogenicity across species.

**Figure S5.**
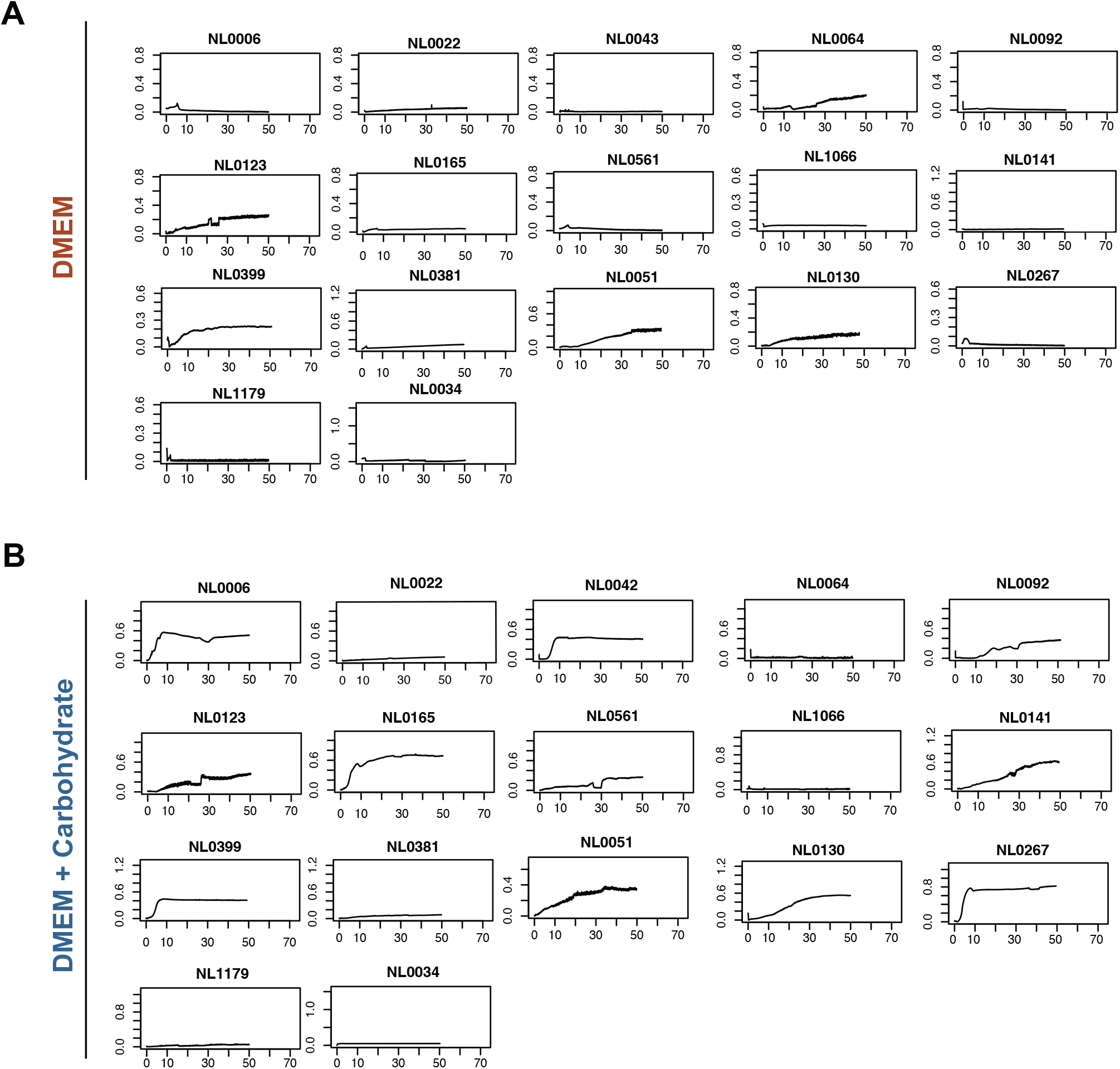
Growth dynamics of in-house bacterial isolates in carbohydrate-supplemented and unsupplemented media. Bacterial strains were cultured in (A) DMEM alone or (B) DMEM supplemented with carbohydrates, and growth was monitored over time. Each panel represents a distinct isolate, with optical density (OD_600_) plotted against time (hours) to assess metabolic adaptability. Strains with a higher proportion of their genome dedicated to carbohydrate-active enzymes (CAZymes) exhibited enhanced growth in the presence of complex carbohydrates.

**Figure S6.**
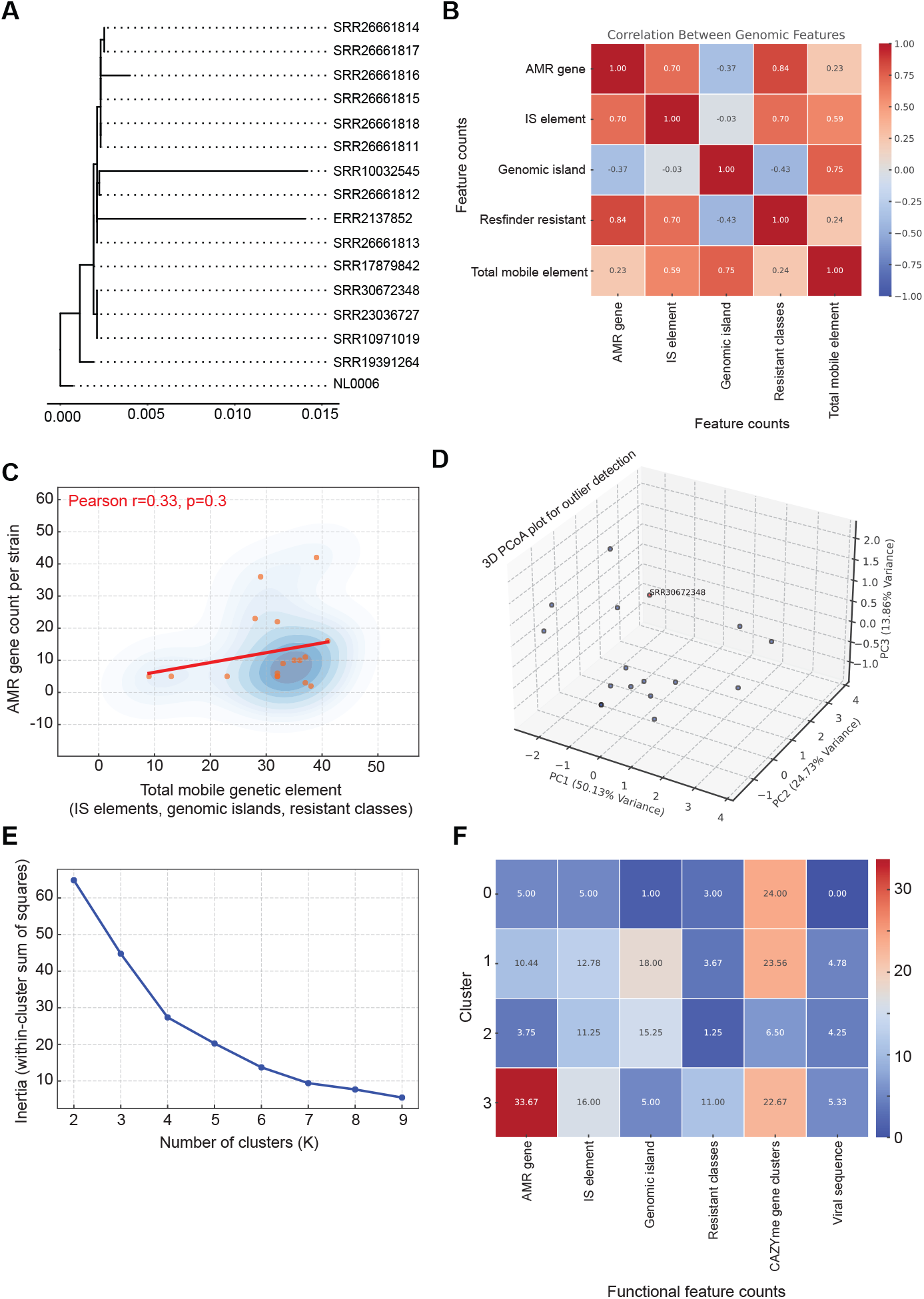
Functional and evolutionary classification of *E. coli* strains. (A) Phylogenetic resolution of *E. coli* strains used in this study with their corresponding Sequence Read Archive (SRA) number. (B) The heatmap illustrates the correlation between key genomic features across *E. coli* strains, highlighting relationships between AMR genes, mobile genetic elements, and genomic islands. (C) Scatter plot shows the relationship between AMR gene load and mobile genetic elements (insertion sequences, genomic islands, and resistant classes). Density contours illustrate the distribution of strains, showing no correlation. (D) A 3D Principal Coordinates Analysis (PCoA) plot identifies functionally distinct outliers. For one strain significantly deviating from the main clusters, the SRA number is plotted, indicating potential adaptive or horizontally transferred genomic elements and highlighting genomic plasticity. (E) The Elbow method plot shows the within-cluster sum of squares (inertia) vs. the number of clusters (K). The inflection point (elbow) at K=4 suggests an optimal cluster number, balancing variance reduction with model complexity. (F) The heatmap shows the functional classification of *E. coli* strains using unsupervised clustering based on key genomic and metabolic features. Strains were grouped into four distinct functional clusters (0-3), highlighting differences in AMR, mobile genetic elements, and metabolic potential including CAZymes.

## Supplementary information

### Supplementary Table 1-12

*Supplementary tables can be downloaded at* https://zenodo.org/records/18479044.

**Table S1. Benchmarking overview**. This table presents key metrics from the benchmarking of *StrainCascade*, summarizing sequencing metadata, assembly statistics, computational resource usage, and sample metadata.

**Table S2. Pairwise comparison of assemblers and annotators**. Performance differences between different assemblers and annotators was performed via paired Wilcoxon signed-rank test for pairwise comparisons. Since each software tool was applied to the same set of samples, the data are paired by sample IDs.

**Table S3. Comparison of assembly metrics across different tools and sequencing platforms**. This table shows a comparative analysis of genome assemblies generated using different assemblers (LJA, SPAdes, Canu, Flye) for various bacterial isolates and provides key assembly statistics, including contig number, genome size, contiguity (N50, N90), and sequencing technology.

**Table S4. Antimicrobial resistance (AMR) genes identified in *E. coli* strains**. The table includes the gene symbol, sequence name, element type, element subtype, resistance class, gene length (bp), reference annotation, and corresponding SRA accession number.

**Table S5. Carbohydrate-active enzyme (CAZyme) information identified in *E. coli* strains**. The table includes the gene type, CAZyme gene cluster-ID, start and end positions, protein family assignment, and corresponding SRA accession number.

**Table S6. Insertion sequences identified in *E. coli* strains**. The table includes the contig tag, transposase cluster, insertion sequence family, sequence length (bp), copy number, element completeness, and corresponding SRA accession number.

**Table S7. The length of pathogenicity island identified in *E*.*coli* strains.**

Table S8. The AMR phenotypes and profiles identified in *E. coli* strains. The table includes the antimicrobial agent, resistance class, predicted genomic AMR phenotype, source file reference, SRA accession number, and taxonomic assignment.

**Table S9. Viral sequences identified in *E. coli* strains**. The table includes viral sequence length (bp), predicted virus group from VirSorter2, source file reference, and corresponding SRA accession number.

**Table S10. Cluster information of Figure 4H**. Mean counts are reported for each category.

**Table S11. Comparison of genome assembly results across heuristic and deterministic runs in *StrainCascade***. This table presents a comparison of genome assembly output files generated by *StrainCascade*, evaluating one heuristic run and three individual deterministic runs per sample. It includes SHA256 hash values to assess bit-identical outcomes, revealing that heuristic results differ from other runs, while repeated deterministic runs produce bit-identical genome assemblies. Please notes that downstream output files cannot be directly compared due to the inclusion of timestamps in their content, limiting the use of hashing to verification of file integrity rather than between-run equivalence.

**Table S12. Computational runtime of *StrainCascade* for bacterial genome analysis across sequencing files**. The table presents the execution time of *StrainCascade* in heuristic and deterministic modes, summarized using key statistical measures

**Table S13. Peak memory usage of *StrainCascade* for bacterial genome analysis across sequencing files**. The table presents the memory requirements of *StrainCascade* in heuristic and deterministic modes, summarized using key statistical measures (GB).

## METHOD DETAILS

### *StrainCascade* setup and execution pipeline

#### Wrapper-based execution for streamlined pipeline workflows

To facilitate seamless execution, *StrainCascade* includes a wrapper script (*StrainCascade_pipeline_wrapper*.*sh*). This wrapper automates the coordination of input files, output directories, and module execution, reducing manual configuration. Users provide key parameters such as input type, sequencing technology, and selected modules. The wrapper dynamically manages directories, sample naming, and Apptainer container calls, ensuring compatibility and modular processing. The modular design allows users to rerun specific stages without duplicating prior steps, optimizing resource usage for high-throughput datasets.

#### Automated pipeline installation and environment setup

Pipeline installation is automated through the *StrainCascade*_installation.sh script, which verifies system prerequisites, including Apptainer installation, directory structure, and disk space. The script offers full or component-specific installation options, guided by user input. It sources utility functions from the *StrainCascade*_installation_utils.sh file to manage checks for required directories and disk availability. This structure simplifies setup across diverse computational environments, minimizing user error during installation.

#### Reproducibility and computational environment

To ensure computational reproducibility, the pipeline employs a fully containerized framework through Apptainer, standardizing system environments across diverse computational setups. All software dependencies employed by modules are encapsulated within different containers, isolating dependencies and eliminating variability caused by differences in software versions or system configurations.

Deterministic execution is enforced through two key strategies: (i) a single-threaded execution mode to avoid variability introduced by parallel processing and (ii) a controlled entropy mechanism based on a fixed entropy source (100 KB of zero bytes). This source is mounted within the container at /*dev/random and /dev/urandom*, minimizing stochastic variability in random number generation during execution.

Although full determinism cannot be achieved for external tools reliant on independent random number generators or system-level entropy, validation tests demonstrated bit-identical outputs across multiple runs for core processes such as genome assembly (Table S11). These measures mitigate non-deterministic behaviour in key analytical steps, supporting consistent, reproducible analyses essential for high-throughput bacterial genome studies.

### Cross-platform dataset selection and composition

A comprehensive dataset of 152 bacterial whole-genome sequencing sourced from the NCBI Sequence Read Archive (SRA) or Yilmaz Lab bacterial isolates from healthy donors as well as patients with a stoma or abdominal surgery (UniBern, Switzerland) were curated to assess the performance of the pipeline (Table S1). Participants were anonymized and collected in an electronic data capture (EDC) system (REDCap) activated for the trial after successfully passing formal quality controls. The Bern EDC system and the database are hosted by the Clinical Trial Unit of Bern University. The collection of luminal content from IBD and non-IBD subjects with stoma was approved by the Bern Cantonal Ethics Commission (BASEC number: 2021-01108 and 2023-00706 with signed informed consent obtained from all participants.

The dataset was designed to capture i) taxonomic diversity, spanning clinically and environmentally significant bacterial genera, and ii) technical diversity, incorporating data from major long-read platforms like PacBio (RS II, Sequel, II/IIe, Revio) and Oxford Nanopore (MinION, GridION). Each platform was optimized for distinct performance parameters, read lengths, and throughput to meet diverse research needs. While complete representation across all platforms was constrained by data availability, the selection strategy ensured comprehensive benchmarking across diverse bacterial taxa and sequencing technologies.

#### Taxonomic Diversity

The dataset spans clinically and environmentally significant bacterial genera, including both pathogenic and commensal species. Key genera represented in the dataset are *Escherichia, Staphylococcus, Streptococcus, Salmonella, Mycobacterium, Klebsiella*, and prominent members of the human gut microbiota, such as *Bacteroides, Bifidobacterium*, and *Akkermansia. Additionally*, the dataset includes Oligo-MM^12^ synthetic bacterial community^31^ and multiple Escherichia coli strains harboring diverse plasmids, enabling the assessment of plasmid detection capabilities.

#### Technical Diversity

The dataset incorporates data from major long-read sequencing platforms, including PacBio (RS II, Sequel, II/IIe, Revio) and Oxford Nanopore (MinION, GridION). These platforms were selected to capture variations in performance parameters, read lengths, and throughput, reflecting the diverse needs of microbial genomics research.

### In-house bacterial isolates and sequencing with PacBio Revio platform

#### Stoma sample collection and bacterial isolation

Stoma content samples were collected from ileostomy and colostomy patients, including individuals with IBD and non-IBD conditions such as cancer. To ensure the viability of strict anaerobes, samples were immediately transferred into sterile 2mL Eppendorf tubes and transported into an anaerobic chamber to prevent oxygen exposure. In parallel, healthy control stool samples were collected, either by mail and promptly frozen upon arrival or in person directly in the lab and cultured freshly.

All culturing was performed under strict anaerobic conditions at 37°C in a Whitley A55 anaerobic workstation. Briefly, single colonies isolated from stoma and fecal samples using rich media were inoculated into 2 mL Eppendorf tubes containing 2 mL of Brain Heart Infusion (BHI, Sigma-Aldrich) liquid medium and incubated for 24 h. Following incubation, 20 μL of the bacterial culture was transferred into a transparent, flat-bottom 96-well plate (Techno Plastic Products), with each well containing 200 μL of growth media. The media tested included Dulbecco’s Modified Eagle Medium/Nutrient Mixture F-12 (DMEM-F12) supplemented with 2.5 mM L-glutamine as the minimal medium, and DMEM-F12 supplemented with 2.5 mM L-glutamine and carbohydrates (Nutricia preOp), containing approximately 12.6 g/L carbohydrates (15% disaccharides, 1% glucose, 9% fructose, 4% maltose, and 70% polysaccharides), to assess which medium facilitated optimal growth in terms of maximum biomass and the time required to reach the mid-log phase. The 96-well plate was then placed in a Stratus Kinetic Microplate Reader (Cerillo), programmed to take measurements at 3-min intervals over 48h. Throughout the measurements, both the plate reader and the 96-well plate were continuously shaken at 500 rpm using Orbital Shaker MS 3 (IKA MS3 basic) to ensure uniform distribution and mixing of the cultures. The maximum biomass and the time required to reach the mid-log phase were modeled in R based on the acquired growth curve

#### Identification and preservation

Single colonies were transferred into 96-well plates containing BHI medium and identified using Bruker Taxonomy and CLOSTRI-TOF (v2.0)^32^ databases on MALDI-TOF MS. After centrifugation and washing with PBS, bacterial pellets were prepared for MALDI-TOF analysis. Colonies of interest were streaked three times for purity before final identification. Once confirmed, bacterial isolates were stored in 2 mL cryotubes with 30% glycerol at -80°C for long-term preservation.

#### DNA extraction and WGS sequencing

Total genomic DNA was extracted from bacterial pellets using a Qiagen MagAttract HMW DNA Kit (Qiagen) following their guidelines for Gram-positive bacterial samples and a Thermo Fisher Scientific Kingfisher Apex robot. The resulting bacterial genomic DNA was assessed for quantity, quality and purity using a Qubit 4.0 fluorometer (Qubit dsDNA HS Assay kit, Thermo Fisher Scientific), an Advanced Analytical FEMTO Pulse instrument (Genomic DNA 165 kb Kit, Agilent) and a Denovix DS-11 UV-Vis spectrophotometer, respectively. Multiplexed SMRTbell libraries were prepared for sequencing on the Revio exactly according to the PacBio guideline entitled: Preparing multiplexed whole genome and amplicon libraries using the HiFi plex prep kit 96. Notably, the gDNA was sheared using a Spex Sample Prep 1600 MiniG 1600 device following the specifications outlined in the technical note from PacBio called ‘High-throughput DNA shearing for long-read microbial’. Thereafter, the sheared gDNA was concentrated and cleaned using 1 x SMRTbell clean-up beads. The samples were then quantified and qualified to be in the range of 5-12Kb using a Qubit 4.0 fluorometer (Qubit dsDNA HS Assay kit, Thermo Fisher Scientific) and an Advanced Analytical FEMTO Pulse instrument (Genomic DNA 165 kb Kit, Agilent), respectively. The remaining steps described earlier included end-repair and A-tailing, followed by the ligation of barcoded overhang adapters and purification of the library using AMPure PB beads along with a nuclease treatment. Subsequently, the libraries were pooled and further purified using AMPure PB beads with a size selection of 3Kb. The concentration and size of the library pool were evaluated again using a ThermoFisher Scientific Qubit 4.0 fluorometer and an Advanced Analytical FEMTO Pulse instrument, respectively, as previously detailed.

Instructions in SMRT Link Sample Setup were followed to prepare the SMRTbell library for sequencing (PacBio SMRT Link v13.1). Shortly, using components from a Revio polymerase kit + cleanup beads bundle (PacBio), the PacBio standard sequencing primer was annealed to the SMRTbell libraries, next the Revio DNA Polymerase was bound, and the polymerase bound complex was bead-based purified. Finally, the Revio sequencing control DNA was diluted and spiked into the complex prior to pipetting onto the thawed Revio sequencing plate (PacBio, PN102-587-400). The Revio deck was setup as directed from the SMRTLink software and included laying out tips, sequencing plates and Revio SMRT Cell trays containing 4 x SMRT cell 25M (PacBio) into their designated locations. The libraries were generally loaded at an on-plate concentration of 300pM using adaptive loading. SMRT sequencing was performed on the Revio controlled by instrument software 13.0.0.212033 or 13.1.0.221972 and with a 30-h movie time. All steps post-bacterial culture harvesting were performed at the Next Generation Sequencing Platform, University of Bern, Switzerland.

### Reads correction and trimming

*StrainCascade* uses Canu^8^ (v2.3) for read correction, trimming, and genome assembly with platform-specific optimizations tailored to both PacBio and Oxford Nanopore Technologies (ONT) sequencing data. The pipeline supports various data types, including raw, corrected, and HiFi reads. For PacBio data, the modes are -*pacbio*, - *pacbio-corr*, and -*pacbio-hifi*, while for ONT data, they include -*nanopore, -nano-corr*, and -*nano-hq*. Input sequencing types determine operations (*-correct, -trim, both or none*), ensuring that pre-processed inputs, such as PacBio HiFi reads, bypass unnecessary steps to optimize performance. This module enforces deterministic execution through a controlled entropy mechanism and offers an optional single-threading mode to minimize run-to-run variability, ensuring reproducibility across different computational environments.

### High-precision deterministic genome assembly

#### Genome assembly

*StrainCascade* integrates four high-performance genome assemblers: La Jolla Assembler (LJA)^11^ (v1.0), SPAdes^9^ (v4.0.0), Canu^8^ (v2.3), and Flye^10^ (v2.9.5-b1801), each optimized for long-read data from both PacBio and ONT. Each assembler is tailored to enhance assembly accuracy for bacterial genomes, with specific platform-aware parameters. To ensure reproducibility, deterministic entropy control is implemented, along with a fully containerized computational environment that mitigates system-level variability leading to reproducible (bit-identical) assembly results (Table S11).

Assemblers requiring genome size estimates, such as Canu^8^ and Flye^10^, utilize an adaptive genome sizing strategy. The pipeline initiates assembly with a default estimate of 4.5 Mb, iteratively refining this based on intermediate results. If prior assembly data are available, these are leveraged to inform subsequent genome size estimates, further improving accuracy. This modular approach, combining adaptive refinement with platform-specific optimizations, delivers reliable assemblies suitable for both *de novo* genome reconstruction and comparative genomic analyses.

#### Assembly assessment

*StrainCascade* evaluates genome quality using complementary tools. QUAST^14^ (v5.3.0) computes assembly contiguity metrics, including N50, L50, and total length. CheckM2^17^ (v1.0.2) estimates genome completeness and contamination through lineage-specific marker genes. Read mapping-based quality control is performed using a dual-mapper approach: NGMLR^15^ (v0.2.7) handles long-read structural variations, while BBMap^16^ (v39.13) manages short-read mapping with *maxindel=100000*. Coverage statistics are generated using pileup.sh script of BBMap, providing per-base coverage metrics alongside overall coverage statistics. This integrated approach, combining assembly metrics, marker gene analysis, and read mapping, ensures a thorough assessment of genome quality and completeness.

#### Assembly refinement

The refinement process follows a sequential optimization approach utilizing MAC2^12^ (v2.1), which implements an adjacency algebraic model with consensus block identification. The pipeline selects an optimal initial assembly—either the best previous assembly or the one with the fewest contigs—and integrates additional assemblies in order of increasing contig counts. A temporal adaptation mechanism limits runtime by filtering out low-quality assemblies after 24 hours if their contig count exceeds three times the median. Circularization is performed with Circlator^13^ (v1.5.5), while sequence accuracy is improved using Arrow algorithm from GenomicConsensus (v2.3.3) for PacBio data (https://github.com/PacificBiosciences/gcpp) and Medaka (v1.11.3) for ONT data (https://github.com/nanoporetech/medaka).

#### Assembly selection

*StrainCascade* completes the assembly process through three stages: initial assembly with LJA, Flye, Canu, and SPAdes; merging using MAC2; and final refinement and circularization with Circlator. Two alternative selection algorithms—*contig* (default) and *continuity*—guide optimization between stages and determine the final assembly based on distinct structural criteria.

The contig algorithm minimizes the number of contigs, favoring assemblies with fewer, larger contigs over those with many smaller ones. It employs a multi-step process, beginning with a plausibility check to eliminate assemblies outside biologically relevant size ranges (>100 Mb or <580 kb). Misassemblies are filtered using a median absolute deviation (MAD) threshold (median + [1.96 × 1.4826 × MAD_size_]), removing extreme outliers. The algorithm retains assemblies with the fewest contigs and, if necessary, performs a length ratio analysis at progressively smaller thresholds (50, 25, 10, 5, 1 kb, and 0 bp) to identify assemblies with the largest proportion of total length covered by contigs. If multiple candidates remain, priority is given to assemblies processed by Circlator or previously favored in earlier *StrainCascade* selections.

The continuity algorithm follows a similar workflow but applies stricter criteria to detect misassemblies. It filters out assemblies with either substantially larger sizes and higher contig counts or those significantly smaller than the median (or <580 kb). This dual-criteria method targets both structural and size-based anomalies. Length ratio analysis mirrors that of the contig algorithm. Final assembly selection considers the fewest contigs, largest maximum contig, overall length, Circlator processing status, and past performance in *StrainCascade* iterations. Equivalent assemblies are documented to ensure reproducibility in each algorithm.

### Taxonomic identification and *de novo* classification

*StrainCascade* integrates taxonomic classification and phylogenetic analysis through Genome Taxonomy Database (GTDB)-Tk^18^ (v2.4.0), utilizing two complementary workflows: classify and *de novo*. This dual approach provides both accurate genome assignment to known taxa and the flexibility to explore novel phylogenetic relationships.

#### Classify workflow

This workflow (classify_wf, with default parameters and --*skip_ani_screen*) serves as the primary method. It assigns query genomes to the Genome Taxonomy Database (GTDB) phylogeny based on a standardized set of 120 bacterial marker genes, ensuring consistent taxonomic placement within the established reference framework.

#### *De novo* workflow

This workflow (*de_novo_wf*, with default parameters, --*bacteria*, and *--outgroup_taxon* set to phylum-level data derived from the classify workflow) supports the construction of custom phylogenetic trees. This workflow is optimized for scalability and can be tailored to include multiple genomes through the *StrainCascade*_GTDB-Tk_de_novo_tree.sh script, enabling broader phylogenetic investigations. While demonstrated with multi-genome input in this publication (Figure S2), it enables researchers to investigate novel lineages and build high-resolution domain-specific phylogenies, providing a powerful tool for *de novo* classification and evolutionary studies.

Taxonomic identification is seamlessly integrated into the interactive HTML analysis report of *StrainCascade*, facilitating accessibility, data visualization, and interpretation. This comprehensive approach enhances both taxonomic precision and exploratory capacity, making it a robust solution for high-throughput bacterial genomic studies.

### Plasmid identification

*StrainCascade* detects plasmid sequences using PlasmidFinder^22^ (v2.1.6), which employs blastn (v2.16.0) to align and identify known plasmid replicons. The analysis is performed with default parameters, including a minimum identity threshold of 90% and a minimum coverage of 60%, ensuring reliable identification.

The workflow begins by retrieving the genome assembly file from the input directory. The identified assembly is temporarily copied to a working directory and processed by plasmidfinder.py. The plasmid reference database is dynamically indexed (INSTALL.py kma_index) to enhance search efficiency. Following analysis, output files (e.g., .json and .txt formats) are renamed with standardized prefixes and transferred to the main output directory. A custom R script (R_process_plasmidfinder.R) processes these results to generate quick-serialization (.qs) files. The final plasmid profiles are integrated into the interactive HTML analysis report, providing users with an intuitive interface for visualization and further exploration of the detected sequences.

### Genome annotation

*StrainCascade* integrates a comprehensive genome annotation pipeline, utilizing Bakta^19^ (v1.8; with --*compliant*), Prokka^20^ (v1.14.6) (with --*compliant, --addgenes, --rfam*, and --*mincontiglen 200*), and MicrobeAnnotator^21^ (v2.0). Each tool is configured with optimized parameters to ensure robust and compliant annotations. For MicrobeAnnotator annotation begins with protein FASTA sequences generated by Bakta^19^ or, alternatively, by Prokka^20^ (using -*m blast* and --*refine*). These tools produce standardized outputs, including GFF3, TSV, and nucleotide/protein FASTA files. Annotation results are integrated and processed using R-based scripts to standardize outputs across tools.

#### Bakta annotation

Bakta^19^ annotation begins by determining taxonomic information based on prior classification results. If available, genus or species-level data is passed to Bakta (--*genus or --species*). The tool operates with default settings (--*compliant, --threads, --locus-tag*) and dynamically indexes its AMRFinderPlus database for antimicrobial resistance annotation. Output files, including .faa, .ffn, and .tsv, are generated and transferred for integration.

#### Prokka annotation

Prokka^20^ annotates genomes with parameters optimized for NCBI compliance. These include --compliant, --addgenes, --rfam, and a minimum contig length of 200 bp (--mincontiglen). The tool assigns locus tags automatically if not provided and supports multi-threaded execution (--cpus). Output formats include .gff, .faa, and .ffn, which are processed for further analyses.

#### MicrobeAnnotator annotation

MicrobeAnnotator^21^ annotates amino acid sequences generated by either Bakta or Prokka. The tool leverages BLAST-based searches (-m blast) against its reference database (--refine). Input .faa files are dynamically managed based on availability and origin (Bakta or Prokka). Key output includes annotated .faa files and KEGG module completeness summaries, supporting downstream functional analysis.

#### Annotation processing

Resulting gene names and products from all tools are standardized and reconciled through a hierarchical string similarity approach implemented in R. Gene names are prioritized based on the following protocol: cross-tool identical names take precedence, gene names are favored over “hypothetical protein” labels, and Bakta names are preferred when discrepancies arise. Name similarity is quantified using the Jaro-Winkler metric, with confidence categories classified as complete (0), high (≤0.15), medium (0.15–0.20), low (0.20–0.45), and none (>0.45). Gene product descriptions are compared using Optimal String Alignment (OSA) distances. Non-hypothetical assignments are prioritized in the order Bakta > Prokka > MicrobeAnnotator. Pairwise OSA distances define consensus confidence categories: complete (all tools agree), high (≤10), medium (10–20), low (20–25), and none (>25).

A weighted scoring system converts these categories to a five-point scale (0-5), assigning 80% weight to gene names and 20% to gene products. Confidence levels are categorized from very low (0-1) to very high (≥4), providing a comprehensive measure of annotation reliability.

#### Functional annotation integration

Enzyme Commission (EC) numbers are consolidated using a hierarchical consensus-based approach to ensure specificity and cross-tool consistency. Predicted EC numbers from each annotation tool are filtered to remove redundancies and grouped based on their hierarchical structure. Redundant entries are collapsed by retaining the most specific variant (e.g., multiple entries such as “1.2.3.4”, “1.2.3.4”, and “1.2.-.-” are consolidated into “1.2.3.4” for a given gene locus). Integration across tools is performed by grouping EC numbers with identical non-dash positions in their four-part classification (e.g., “1.1.1.1” and “1.1.-.-” are grouped together). EC numbers are scored based on their level of completeness (1–4), with the most specific and compatible entry selected as the primary assignment (*EC_number_SC_best*). Alternative assignments are retained with associated confidence scores (*EC_number_SC_all*).

Clusters of Orthologous Groups (COG) annotations are integrated from Bakta and Prokka. If a COG prediction is available from only one tool, it is retained as the final assignment. In cases where both tools provide predictions, the assignment from Bakta is prioritized due to its broader database coverage. Functional categories linked to the COG numbers are also derived from Bakta and included in the unified output (COG_number_SC, COG_category_SC).

KEGG orthology (K numbers) annotations are reconciled from Bakta and MicrobeAnnotator using a confidence-based integration framework. When predictions from both tools overlap, Bakta assignments are prioritized to produce a unified set of annotations *(K_number_SC*), enhancing both accuracy and consistency for downstream functional analyses.

### Genomic islands identification

The pipeline identifies genomic islands (GIs) using IslandPath-DIMOB (v1.0.6) pipeline^23^, which integrates sequence composition analysis and mobility gene detection. The method targets genomic regions with significant dinucleotide composition deviations and clusters of eight or more genes, along with the presence of at least one mobility-associated gene. Input files in GenBank format (e.g., .gbff from Bakta or .gbk from Prokka) are preprocessed and standardized to avoid file name conflicts before analysis.

The *Dimob* module generates .gff3 output files containing the identified genomic islands, which are then incorporated into the pipeline for further integration and visualization. Results are processed by an R script (R_process_islandpath.R), which consolidates both the GI features and the corresponding nucleotide sequences for seamless data interpretation into the interactive HTML analysis report. This hierarchical approach is essential for investigating horizontal gene transfer and genome evolution in bacterial populations.

### Metabolic pathway analysis

Metabolic pathway analysis using dbCAN3^33^ (v4.1.4) for carbohydrate-active enzyme (CAZyme) detection and MicrobeAnnotator^21^ (v2.0) for KEGG pathway evaluation are integrated to deliver robust insights into both carbohydrate metabolism and broader metabolic networks in bacterial genomes, respectively. CAZyme identification utilizes multiple search tools (*--tools all*), including HMMER, DIAMOND, and Hotpep, with default parameters for threshold and coverage settings. The analysis encompasses subfamily-level substrate predictions and the identification of CAZyme gene clusters (CGCs) through clustering (--*cluster 1*) and CGC detection (--*cgc_sig_genes all, --cgc_dis 2*). Output files such as overview.txt and cgc_standard.out summarize the identified enzymes and gene clusters. These results are further processed using an R script (R_process_dbcan3.R) to standardize outputs. MicrobeAnnotator assesses metabolic potential by evaluating KEGG module completeness through targeted database searches, providing a detailed overview of functional gene modules within metabolic pathways. The results from both analyses are integrated into an interactive HTML report for visualization and exploration, with quick-serialization files (.qs) generated to support streamlined downstream analyses. This comprehensive approach delivers robust insights into both carbohydrate metabolism and broader metabolic networks in bacterial genomes.

### Antimicrobial resistance gene analysis

Antimicrobial resistance (AMR) genes are identified using two complementary tools: i) ResFinder^24^ (v4.6.0) and ii) AMRFinderPlus^25^ (v4.0.3). ResFinder detects acquired resistance genes and chromosomal mutations, utilizing parameters optimized for coverage and thresholds set at 60% for minimum gene length and 80% for sequence identity. For mutation detection, additional options (--*point, --l_p 0*.*6, --t_p 0*.*8*) are applied. Key options include *--acquired, --point, --ignore_missing_species, -acq, -d*, and *-u*. AMRFinderPlus expands the analysis by identifying resistance genes and mutations using the NCBI AMRFinderPlus database, employing options such as *--plus and -- mutation_all*. Both tools generate detailed outputs on resistance phenotypes and mechanisms, which are integrated into an interactive HTML report for enhanced visualization and exploration.

### Characterization of genome plasticity

Genome plasticity is characterized through the precise identification of defense systems, mobile elements, and viral sequences using CRISPRCasFinder^26^, ISEScan^34^, VirSorter2^27^ (v2.2.4) and DeepVirFinder^28^ (v1.0) all implemented in *StrainCascade* with default parameters. The combined results provide a comprehensive profile of viral elements, defense mechanisms, and mobile genetic entities, offering insights into horizontal gene transfer and genome adaptation.

Genome plasticity is characterized through the precise and reproducible identification of defense systems and mobile elements using the CRISPRCasFinder^26^ and ISEScan^34^ pipelines. Viral sequences are detected using VirSorter2^27^ (v2.2.4) and DeepVirFinder^28^ (v1.0) both implemented in *StrainCascade* and executed with default parameters to ensure robust detection. By synthesizing the results obtained from these analyses, a comprehensive profile of viral elements, defense mechanisms, and mobile genetic entities within the dataset may be produced, offering novel insights into horizontal gene transfer and genomic plasticity.

#### CRISPR-Cas systems detection

CRISPRCasFinder^26^ detects CRISPR-Cas systems by combining CRISPR array detection with Cas protein identification. It employs structural similarity-based analysis (-*so*) and multi-threaded execution (*-cpuM*) to enhance performance. The input assembly is preprocessed, and temporary files are generated during processing. The output includes various file formats, such as .gff, .tsv, and .json, which summarize CRISPR arrays, Cas proteins, and their genomic contexts. These results are subsequently standardized and transferred for integration into the analysis pipeline.

#### Insertion sequence (IS) elements

They are detected using ISEScan^34^, which identifies both IS elements and their associated open reading frames (ORFs). The analysis leverages multi-threading (--*nthread*) for efficiency and outputs annotations in .gff, .tsv, and .fna formats. The detected IS elements and ORFs are further processed by an R script (R_process_isescan.R) to consolidate data and generate quick-serialization outputs.

#### Viral sequence detection

Viral sequences are detected using VirSorter2^27^ (v2.2.4) and DeepVirFinder (v1.0)^28^ .VirSorter2 applies Hidden Markov Models (HMMs) to predict viral sequences based on reference databases, with parameters configured to set a minimum sequence length of 200 bp (--*min-length 200*). Output files, including .fa and .tsv reports, provide detailed information on viral boundaries and scores, which are processed and standardized by the R script R_process_virsorter2.R. DeepVirFinder^28^ leverages a deep learning framework for reference-free viral prediction, using CPU-based optimization (THEANO_FLAGS) and a minimum input length of 200 bp (*-l 200*). The results, saved as prediction files (dvfpred.txt), are integrated into the pipeline’s functional analysis.

The outputs, including CRISPR-Cas systems, IS elements, and viral sequences, are consolidated into an interactive HTML report, enabling comprehensive visualization and exploration of these genomic features.

## QUANTIFICATION AND STATISTICAL ANALYSIS

### Non-parametric comparisons of assembly and annotation performance

Differences in performance across genome assemblers and annotation tools were first assessed using non-parametric frequentist approaches. For assembly performance (number of contigs, N50) and annotation performance (number of CDS, EC number completeness), we applied paired Wilcoxon signed-rank tests. Pairing was based on sample IDs, as each tool was applied to the same set of samples. Significance was determined using a threshold of α = 0.05, with Bonferroni correction for multiple testing.

### Hierarchical Bayesian modeling

Hierarchical Bayesian models was used ro quantify tool-specific performance more robustly. A negative binomial regression model was used for over-dispersed count data (e.g., number of contigs), while Beta regression models were employed for bounded proportional data (e.g., N50 ratios and EC completeness). Random effects for tools (assembler or annotator) and sample IDs were included to account for tool-specific effects and inter-sample variability.

All models were fitted using four Markov chains run for 12,000 iterations (6,000 warmup), with an adaptation delta of 0.9999 and a maximum tree depth of 20 to ensure stable convergence. Diagnostcs included the Gelman–Rubin statstc 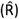 and effective sample size. Weakly informative normal priors (μ = 0, σ = 1) were placed on the logit scale for proportion-based models and the log scale for count models. This structure supports estimation of tool-specific performance while respecting the nested design of the experiment.

### Posterior analysis and reporting

Posterior distributions were analyzed using two complementary approaches. First, probabilities of superiority (P(best)) were derived by comparing tools across posterior draws, identifying the likelihood that a given tool outperformed other. For assembly contigs, lower values were considered optimal; for N50 ratio and EC completeness, higher values were preferred. Second, we assessed the magnitude and uncertainty of effects by computing deviations from the global mean on the probability scale using logistic transformations. Results are reported as mean posterior estimates with 95% credible intervals.

All Bayesian analyses were conducted in R (v2.22.0) using the brms package with Stan as the backend for Bayesian computation, and a fixed random seed (123) to ensure reproducibility.

### Computing resource

All computations were performed on the High-Performance Computing (HPC) cluster of the University of Bern. The cluster operates on Rocky Linux 9.4 (Blue Onyx) with kernel version 5.14.0-427.40.1.el9_4.x86_64 and an x86_64 architecture. Jobs based on node availability, utilizing either an AMD EPYC 7742 (64-core) Processor or an AMD EPYC 9654 (96-core) Processor. All sequencing files were processed twice with *StrainCascade*. Once in heuristic mode with 32 CPUs allocated and once in deterministic mode with 1 CPU allocated.

*StrainCascade* runtime varies based on processing mode and dataset complexity. In heuristic mode, optimized for speed, the median runtime is 2 hours 32 minutes (1h 51m 27s for PacBio HiFi by Revio), with a range from 51 minutes to 68 hours 49 minutes (Table S12). The deterministic mode, prioritizing accuracy, has a median runtime of 9 hours and 42 minutes, ranging from 1 hour to 46 minutes and 53 hours to 43 minutes. The broader range reflects variations in sequencing depth, genome complexity, and computational resources.

*StrainCascade* memory requirements remain consistent across processing modes. Both heuristic and deterministic modes exhibit identical median peak memory usage (71 GB) with similar interquartile ranges (58.5-74 GB and 57.25-74 GB, respectively), ranging from 20-88 GB in heuristic mode and 10-81 GB in deterministic mode (Table S13). This indicates that processing mode selection has minimal impact on memory consumption compared to the inherent variability between genomic samples.

## Notes

https://github.com/SBUJordi/StrainCascade

https://sbujordi.github.io/StrainCascade_documentation/

https://zenodo.org/records/14921436

### Competing Interest Statement

The authors have declared no competing interest.

## References

1. Hou, K., Wu, Z.X., Chen, X.Y., Wang, J.Q., Zhang, D., Xiao, C., Zhu, D., Koya, J.B., Wei, L., Li, J., and Chen, Z.S. (2022). Microbiota in health and diseases. Signal Transduct Target Ther 7, 135. 10.1038/s41392-022-00974-4.

2. Agustinho, D.P., Fu, Y., Menon, V.K., Metcalf, G.A., Treangen, T.J., and Sedlazeck, F.J. (2024). Unveiling microbial diversity: harnessing long-read sequencing technology. Nature methods 21, 954–966. 10.1038/s41592-024-02262-1.

3. Quijada, N.M., Rodriguez-Lazaro, D., Eiros, J.M., and Hernandez, M. (2019). TORMES: an automated pipeline for whole bacterial genome analysis. Bioinformatics 35, 4207–4212. 10.1093/bioinformatics/btz220.

4. Petit, R.A., 3rd, and Read, T.D. (2020). Bactopia: a Flexible Pipeline for Complete Analysis of Bacterial Genomes. mSystems 5. 10.1128/mSystems.00190-20.

5. Schwengers, O., Hoek, A., Fritzenwanker, M., Falgenhauer, L., Hain, T., Chakraborty, T., and Goesmann, A. (2020). ASA3P: An automatic and scalable pipeline for the assembly, annotation and higher-level analysis of closely related bacterial isolates. PLoS computational biology 16, e1007134. 10.1371/journal.pcbi.1007134.

6. Zhou, Z., Alikhan, N.F., Mohamed, K., Fan, Y., Agama Study, G., and Achtman, M. (2020). The EnteroBase user’s guide, with case studies on Salmonella transmissions, Yersinia pestis phylogeny, and Escherichia core genomic diversity. Genome Res 30, 138–152. 10.1101/gr.251678.119.

7. Espinosa, E., Bautista, R., Larrosa, R., and Plata, O. (2024). Advancements in long-read genome sequencing technologies and algorithms. Genomics 116, 110842. 10.1016/j.ygeno.2024.110842.

8. Koren, S., Walenz, B.P., Berlin, K., Miller, J.R., Bergman, N.H., and Phillippy, A.M. (2017). Canu: scalable and accurate long-read assembly via adaptive k-mer weighting and repeat separation. Genome Res 27, 722–736. 10.1101/gr.215087.116.

9. Prjibelski, A., Antipov, D., Meleshko, D., Lapidus, A., and Korobeynikov, A. (2020). Using SPAdes De novo Assembler. Curr Protoc Bioinformatics 70, e102. 10.1002/cpbi.102.

10. Kolmogorov, M., Yuan, J., Lin, Y., and Pevzner, P.A. (2019). Assembly of long, error-prone reads using repeat graphs. Nat Biotechnol 37, 540–546. 10.1038/s41587-019-0072-8.

11. Bankevich, A., Bzikadze, A.V., Kolmogorov, M., Antipov, D., and Pevzner, P.A. (2022). Multiplex de Bruijn graphs enable genome assembly from long, high-fidelity reads. Nat Biotechnol 40, 1075–1081. 10.1038/s41587-022-01220-6.

12. Tang, L., Li, M., Wu, F.X., Pan, Y., and Wang, J. (2019). MAC: Merging Assemblies by Using Adjacency Algebraic Model and Classification. Front Genet 10, 1396. 10.3389/fgene.2019.01396.

13. Hunt, M., Silva, N.D., Otto, T.D., Parkhill, J., Keane, J.A., and Harris, S.R. (2015). Circlator: automated circularization of genome assemblies using long sequencing reads. Genome Biol 16, 294. 10.1186/s13059-015-0849-0.

14. Mikheenko, A., Prjibelski, A., Saveliev, V., Antipov, D., and Gurevich, A. (2018). Versatile genome assembly evaluation with QUAST-LG. Bioinformatics 34, i142–i150. 10.1093/bioinformatics/bty266.

15. Sedlazeck, F.J., Rescheneder, P., Smolka, M., Fang, H., Nattestad, M., von Haeseler, A., and Schatz, M.C. (2018). Accurate detection of complex structural variations using single-molecule sequencing. Nature methods 15, 461–468. 10.1038/s41592-018-0001-7.

16. Wang, X., Jowsey, W.J., Cheung, C.Y., Smart, C.J., Klaus, H.R., Seeto, N.E., Waller, N.J., Chrisp, M.T., Peterson, A.L., Ofori-Anyinam, B., et al. (2024). Whole genome CRISPRi screening identifies druggable vulnerabilities in an isoniazid resistant strain of Mycobacterium tuberculosis. Nature communications 15, 9791. 10.1038/s41467-024-54072-w.

17. Chklovski, A., Parks, D.H., Woodcroft, B.J., and Tyson, G.W. (2023). CheckM2: a rapid, scalable and accurate tool for assessing microbial genome quality using machine learning. Nature methods 20, 1203–1212. 10.1038/s41592-023-01940-w.

18. Chaumeil, P.A., Mussig, A.J., Hugenholtz, P., and Parks, D.H. (2019). GTDB-Tk: a toolkit to classify genomes with the Genome Taxonomy Database. Bioinformatics 36, 1925–1927. 10.1093/bioinformatics/btz848.

19. Schwengers, O., Jelonek, L., Dieckmann, M.A., Beyvers, S., Blom, J., and Goesmann, A. (2021). Bakta: rapid and standardized annotation of bacterial genomes via alignment-free sequence identification. Microb Genom 7. 10.1099/mgen.0.000685.

20. Seemann, T. (2014). Prokka: rapid prokaryotic genome annotation. Bioinformatics 30, 2068–2069. 10.1093/bioinformatics/btu153.

21. Ruiz-Perez, C.A., Conrad, R.E., and Konstantinidis, K.T. (2021). MicrobeAnnotator: a user-friendly, comprehensive functional annotation pipeline for microbial genomes. BMC Bioinformatics 22, 11. 10.1186/s12859-020-03940-5.

22. Carattoli, A., Zankari, E., Garcia-Fernandez, A., Voldby Larsen, M., Lund, O., Villa, L., Moller Aarestrup, F., and Hasman, H. (2014). In silico detection and typing of plasmids using PlasmidFinder and plasmid multilocus sequence typing. Antimicrobial agents and chemotherapy 58, 3895–3903. 10.1128/AAC.02412-14.

23. Bertelli, C., and Brinkman, F.S.L. (2018). Improved genomic island predictions with IslandPath-DIMOB. Bioinformatics 34, 2161–2167. 10.1093/bioinformatics/bty095.

24. Florensa, A.F., Kaas, R.S., Clausen, P., Aytan-Aktug, D., and Aarestrup, F.M. (2022). ResFinder - an open online resource for identification of antimicrobial resistance genes in next-generation sequencing data and prediction of phenotypes from genotypes. Microb Genom 8. 10.1099/mgen.0.000748.

25. Feldgarden, M., Brover, V., Gonzalez-Escalona, N., Frye, J.G., Haendiges, J., Haft, D.H., Hoffmann, M., Pettengill, J.B., PrASAd, A.B., Tillman, G.E., et al. (2021). AMRFinderPlus and the Reference Gene Catalog facilitate examination of the genomic links among antimicrobial resistance, stress response, and virulence. Scientific reports 11, 12728. 10.1038/s41598-021-91456-0.

26. Grissa, I., Vergnaud, G., and Pourcel, C. (2007). CRISPRFinder: a web tool to identify clustered regularly interspaced short palindromic repeats. Nucleic Acids Res 35, W52–57. 10.1093/nar/gkm360.

27. Guo, J., Bolduc, B., Zayed, A.A., Varsani, A., Dominguez-Huerta, G., Delmont, T.O., Pratama, A.A., Gazitua, M.C., Vik, D., Sullivan, M.B., and Roux, S. (2021). VirSorter2: a multi-classifier, expert-guided approach to detect diverse DNA and RNA viruses. Microbiome 9, 37. 10.1186/s40168-020-00990-y.

28. Ren, J., Song, K., Deng, C., Ahlgren, N.A., Fuhrman, J.A., Li, Y., Xie, X., Poplin, R., and Sun, F. (2020). Identifying viruses from metagenomic data using deep learning. Quant Biol 8, 64–77. 10.1007/s40484-019-0187-4.

29. Jung, J.M., Rahman, A., Schiffer, A.M., and Weisberg, A.J. (2024). Beav: a bacterial genome and mobile element annotation pipeline. mSphere 9, e0020924. 10.1128/msphere.00209-24.

30. Wick, R.R., Judd, L.M., Cerdeira, L.T., Hawkey, J., Meric, G., Vezina, B., Wyres, K.L., and Holt, K.E. (2021). Trycycler: consensus long-read assemblies for bacterial genomes. Genome Biol 22, 266. 10.1186/s13059-021-02483-z.

31. Brugiroux, S., Beutler, M., Pfann, C., Garzetti, D., Ruscheweyh, H.J., Ring, D., Diehl, M., Herp, S., Lotscher, Y., Hussain, S., et al. (2016). Genome-guided design of a defined mouse microbiota that confers colonization resistance against Salmonella enterica serovar Typhimurium. Nat Microbiol 2, 16215. 10.1038/nmicrobiol.2016.215.

32. ASAre, P.T., Lee, C.H., Hurlimann, V., Teo, Y., Cuenod, A., Akduman, N., Gekeler, C., Afrizal, A., Corthesy, M., Kohout, C., et al. (2023). A MALDI-TOF MS library for rapid identification of human commensal gut bacteria from the class Clostridia. Front Microbiol 14, 1104707. 10.3389/fmicb.2023.1104707.

33. Zheng, J., Ge, Q., Yan, Y., Zhang, X., Huang, L., and Yin, Y. (2023). dbCAN3: automated carbohydrate-active enzyme and substrate annotation. Nucleic Acids Res 51, W115–W121. 10.1093/nar/gkad328.

34. Xie, Z., and Tang, H. (2017). ISEScan: automated identification of insertion sequence elements in prokaryotic genomes. Bioinformatics 33, 3340–3347. 10.1093/bioinformatics/btx433.

